# What does it take to learn the rules of RNA base pairing? A lot less than you may think

**DOI:** 10.1101/2025.07.31.668042

**Authors:** Jayanth S. Pratap, Ryan K. Krueger, Elena Rivas

## Abstract

Amidst the fast-developing trend of RNA large language models with millions of parameters, we asked what would be the minimum required to rediscover the rules of RNA canonical base pairing, mainly the Watson-Crick-Franklin A:U, G:C and the wobble G:U base pairs (the secondary structure). Here, we conclude that it does not require much at all. It does not require knowing secondary structures; it does not require aligning the sequences; and it does not require many parameters. We selected a probabilistic model of palindromes (a stochastic context-free grammar or SCFG) with a total of just 21 parameters. Using standard deep learning techniques, we estimate its parameters by implementing the generative process in an automatic differentiation (autodiff) framework and applying stochastic gradient descent (SGD). We define and minimize a loss function that does not use any structural or alignment information. Trained on as few as fifty RNA sequences, the rules of RNA base pairing emerge after only a few iterations of SGD. Crucially, the sole inputs are RNA sequences. When optimizing for sequences corresponding to structured RNAs, SGD also yields the rules of RNA base-pair aggregation into helices. Trained on shuffled sequences, the system optimizes by avoiding base pairing altogether. Trained on messenger RNAs, it reveals interactions that are different from those of structural RNAs, and specific to each mRNA. Our results show that the emergence of canonical base-pairing can be attributed to sequence-level signals that are robust and detectable even without labeled structures or alignments, and with very few parameters. Autodiff algorithms for probabilistic models, such as, but not restricted to SCFGs, have significant potential as they allow these models to be incorporated into end-to-end RNA deep learning methods for discerning transcripts of different functionalities.

**Availability:** **rivaslab.org, https://github.com/EddyRivasLab/R-scape/tree/master/python/d-SCFG**

## Introduction

### There are many large language models, but what is the minimum sufficient to infer RNA base pairing?

RNA language models used for RNA secondary and tertiary structure prediction rely on millions of parameters. For instance, RiNALMo [17], one of the largest RNA models so far, uses 650 million parameters. Similarly, other deep learning methods to infer RNA secondary structure such as MXFOLD2 [23] or UFold [9] use hundreds of thousands of parameters and train on many thousands of structures. Some methods such as SPOT-RNA2 [25] also include alignments. Deep learning methods are often celebrated for learning RNA base pairing [24], but what would the performance of much simpler models trained with similar methods look like?

Here, we sought to test the minimal set of requirements to infer RNA base pairing. In particular, we wanted to test three aspects: (1) whether there is a need to train on known structures; (2) whether in the absence of structures, it is necessary to train on alignments which carry information about covariation between base-paired position [21]; and (3) the size of a minimal model that could learn the rules of RNA base pairing.

We have approached these questions by looking at probabilistic models of RNA base pairing, named stochastic context-free grammars (SCFGs). SCFGs, first established in the context of natural languages, are suitable for modeling nested pairwise interactions like those observed in RNA structures. The probabilistic parameters of an SCFG are easy to train from structural data by maximum likelihood. Here, we concentrate on several small and simple SCFG designs that have been shown to have prediction accuracies near the performance of standard thermodynamic models that depend on thousands of parameters [6].

The training of parameters in modern deep learning is typically performed via stochastic gradient descent (SGD), an optimization method that requires calculating gradients, and modifying the parameters with quantities opposite to their gradients. These gradients are computed using automatic differentiation (autodiff), a technique that takes a function and automatically constructs a procedure to compute its derivatives (the gradient values). In practice, this is often implemented via backpropagation, an autodiff algorithm that is particularly memory efficient for large neural networks. We implemented minimal SCFGs for RNA secondary structure in JAX [1], a state-of-the-art automatic differentiation framework, to enable efficient gradient calculations with respect to SCFG parameters. By applying SGD, we systematically explored the conditions under which the optimal parameters trained in the absence of prior structural information or alignments are able to reproduce the rules of RNA base pairing, that is, the Watson-Crick-Franklin A:U, G:C, and the wobble G:U base pairs (WCF base pairs) that arrange into helices forming the secondary structure.

There are other optimization methods to train from unlabeled (unstructured) data such as the Baum-Welsh expectation-maximization algorithm (EM) [18]. EM has been successfully applied to train SCFGs for specific structural RNA families (covariance models) without using any prior structural or alignment information [7]. Our question is related but different, as we try to infer the elements common to all RNA structures with the minimal SCFG able to model–not just one structure per model (for homology searching), but all possible RNA structures with one model (for secondary structure prediction).

Our approach of training SCFGs by SGD has not been explored before. The advantage of SGD over EM is that SGD only requires implementing one algorithm in an automatic differentiation framework (JAX), enabling the automatic calculation of the gradients and loss. In addition, JAX enables compilation to hardware accelerators such as GPUs and TPUs. This gives us the versatility to test many different SCFGs using standard deep neural network (DNN) training methods, which also facilitates a direct integration of these SCFGs into other more complex DNNs, such as those investigating RNA 3D structure.

## Results

### A minimal SCFG for RNA base pairing

SCFGs are probabilistic models designed to describe pairwise correlations such as palindromes or RNA base pairs forming helices [6, 22]. SCFGs as complex as standard thermodynamic models (that include thousands of parameters) produce comparable results using algorithms of similar complexity [22], but lightweight SCFGs with just a handful of parameters have also been shown to perform comparably to standard thermodynamic models [6].

For this minimal experiment, we selected the G6 grammar [6], first introduced in the method PFOLD [11] and described in Figure 1. The G6 grammar includes only a total of 21 independent parameters, and consists of three non-terminals. For each non-terminal (*S, L, F*), there is a discrete probability distribution (*T*_*S*_, *T*_*L*_, *T*_*F*_), which assigns a probability to each of the non-terminal’s rules. Figure 1a shows how the complete set of probability parameters for the G6 SCFG can be described with three Bernoulli probabilities, together with two emission probability distributions as,

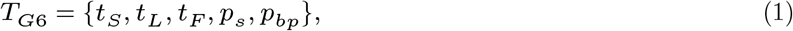

where, *t*_*S*_, *t*_*L*_, *t*_*F*_ are Bernoulli probabilities for rule decisions at each nonterminal *S, L, F* ; *p*_*s*_(*a*) is the emission probability for single nucleotide *a*; and *p*_*bp*_(*a, b*) is the emission probability for base pair (*a, b*), where *a, b* ∈ {*A, C, G, U*} and ∑_*a*_ *p*_*s*_(*a*) = 1, ∑_*ab*_ *p*_*bp*_(*a, b*) = 1.

**Figure 1:**
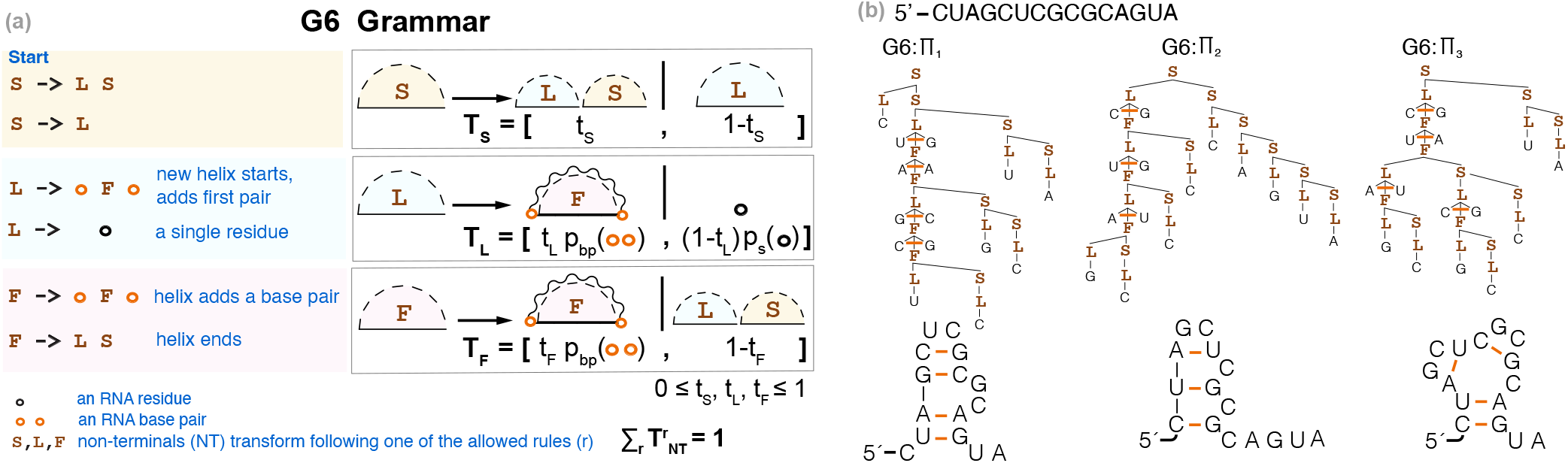
The G6 SCFG models nested helices of RNA base pairs. **(a)** Description of the G6 grammar. Each non-terminal, *L, S*, and *F*, has a discrete probability distribution, *T*_*S*_, *T*_*L*_, *T*_*F*_, describing the probabilities over their rules. S is the start non-terminal. **(b)** For a toy sequence of length 15, we show examples of three different possible secondary structures and how those get parsed and assigned a probability by the G6 grammar. See Eq. 1 for parameter definitions.

The base pair probabilities, *p*_*bp*_, could be made more complex by distinguishing between those at the start of a helix (used with *L*) from those inside a helix (used with *F*); the latter could also be made conditional on the previous base pair (usually referred to as base pair stacking). Here, we wanted to start with the simplest possible implementation using only one single base pair probability and disregarding stacking (Figure 1a). This simplified G6 parameterization (Eq. 1) has been shown to be enough for our purpose of identifying WCF base pairing when trained using sequences and structures [6].

While the focus is on canonical (WCF and wobble) base pairs, notice that the base pair probabilities could in principle capture non-canonical base pairs, if those were prominent in structural RNAs. And while SCFGs cannot capture pseudoknots, by integrating to all possible structures, all helices involved in possible pseudoknots would be taken into account.

### Parameter optimization from sequence alone by integrating over all possible structures

SCFGs are usually trained by maximum likelihood which requires using datasets of known structured RNAs with known secondary structures. Alternatively, we show that SCFGs can be trained using only RNA sequences by optimizing the probability of the sequence, given the model, without needing structure annotations.

For a given sequence (*seq*) and a possible structure for that sequence (*π*_*seq*_), there is one (and only one, if the grammar is unambiguous) parse and one probability score *P* (*seq, π*_*seq*_ ∣ *G, T*_*G*_) associated with that sequence/structure pair. The probability of a given sequence is then calculated by summing the contributions of all possible structures, *π*_*seq*_, that are consistent with that sequence:

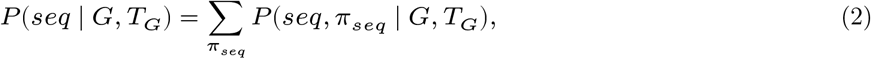

where *G* is the grammar, *T*_*G*_ is its set of parameters, and the sum is over all secondary structures *π*_*seq*_ compatible with *seq*.

Figure 1b shows examples of different structures associated with a given sequence, and how those get parsed by the G6 SCFG, which is unambiguous.

While the direct enumeration of the contribution of all possible structures is highly inefficient, there is a dynamic programming algorithm, named the Inside algorithm, that calculates the probability of a sequence by summing the contributions of all possible structures in time polynomial in the length of the sequence. The Inside algorithm is analogous to the McCaskill algorithm [16] used by the standard thermodynamic models of RNA folding (such as Mfold [29], ViennaRNA [14], or RNAstructure [19]) to calculate the partition function [20, 8].

For instance, for a sequence of length *n*, (*s*_1_ … *s*_*n*_), the G6 Inside algorithm calculates *n* × *n* matrices **I**_*S*_, **I**_*L*_, **I**_*F*_, such that for each *i* ≤ *j* = {1, …, *n*},

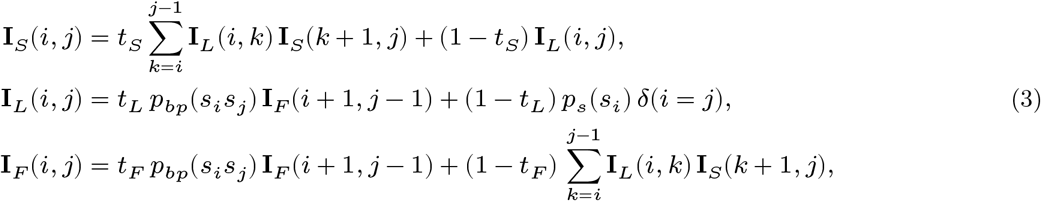

where **I**_*S*_(*i, j*), **I**_*L*_(*i, j*), **I**_*F*_ (*i, j*) are Inside scores for substring *s*_*i*_ … *s*_*j*_ under *S, L*, and *F* non-terminals respectively. The polynomial time complexity of the G6 Inside algorithm is 𝒪(*n*^3^), where *n* is the sequence length.

For any SCFG, the Inside probability for a sequence is the total probability summed over all possible parses (secondary structures) from the start non-terminal *S*. We denote,

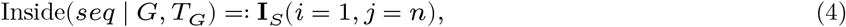

so that the probability of observing sequence *seq* under grammar *G* and parameters *T*_*G*_ is given by

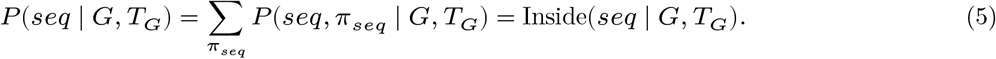

We have produced differentiable implementations of the Inside algorithm for the grammars tested here using the Python library JAX, a popular automatic differentiation library for scientific computing [1]. Our implementation enables backpropagation through the Inside algorithm, and therefore gradient-based parameter updates. These are formally equivalent to differentiable implementations of the McCaskill algorithm for computing the partition function for thermodynamic models [15, 13, 12].

Using these differentiable implementations of the Inside algorithm for a given SCFG, we perform SGD to optimize the probabilistic parameters. Given a training set of RNA sequences with no structure annotation, the optimal parameters are obtained by

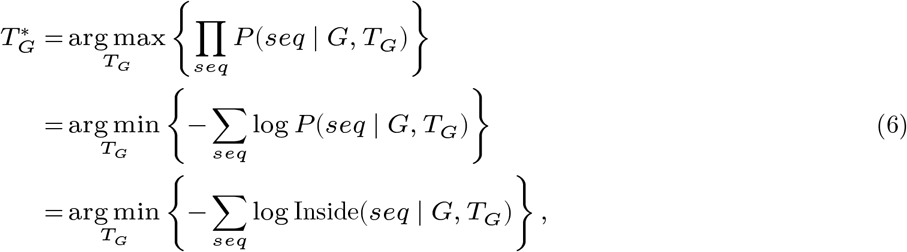

where the loss to be minimized is the negative log-likelihood, *i*.*e*., minus the sum over all training sequences of the log-probability assigned by the current SCFG. The sum is over all sequences in the batch; the Inside algorithm automatically marginalizes over all possible structures for each sequence. At each epoch, SGD is applied to a batch of 10 sequences selected at random from the training set.

In particular for G6, automatic differentiation using RNA sequences alone with no corresponding structures permits numerical parameter optimization to obtain

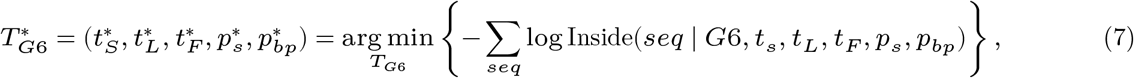

for the Inside algorithm described in Eq. 3, with the parameters as in Eq. 1.

### RNA base pairing rules emerge rather quickly

Figure 2 describes the training of the G6 grammar on a collection of 50 RNaseP RNA sequences, selected at random from a database of 225 RNaseP sequences introduced before [6] from the Ribonuclease P database [2]. We observe that the WCF base pairing rules begin to emerge in the pair probability distribution after only a few epochs of training. Notably, no structural information is provided during training. In Supplemental Figure S1, we report that even when reducing the training set to 25 RNaseP sequences, the resulting model still produces reasonably accurate base pairing probabilities, although the loss becomes more unstable.

**Figure 2:**
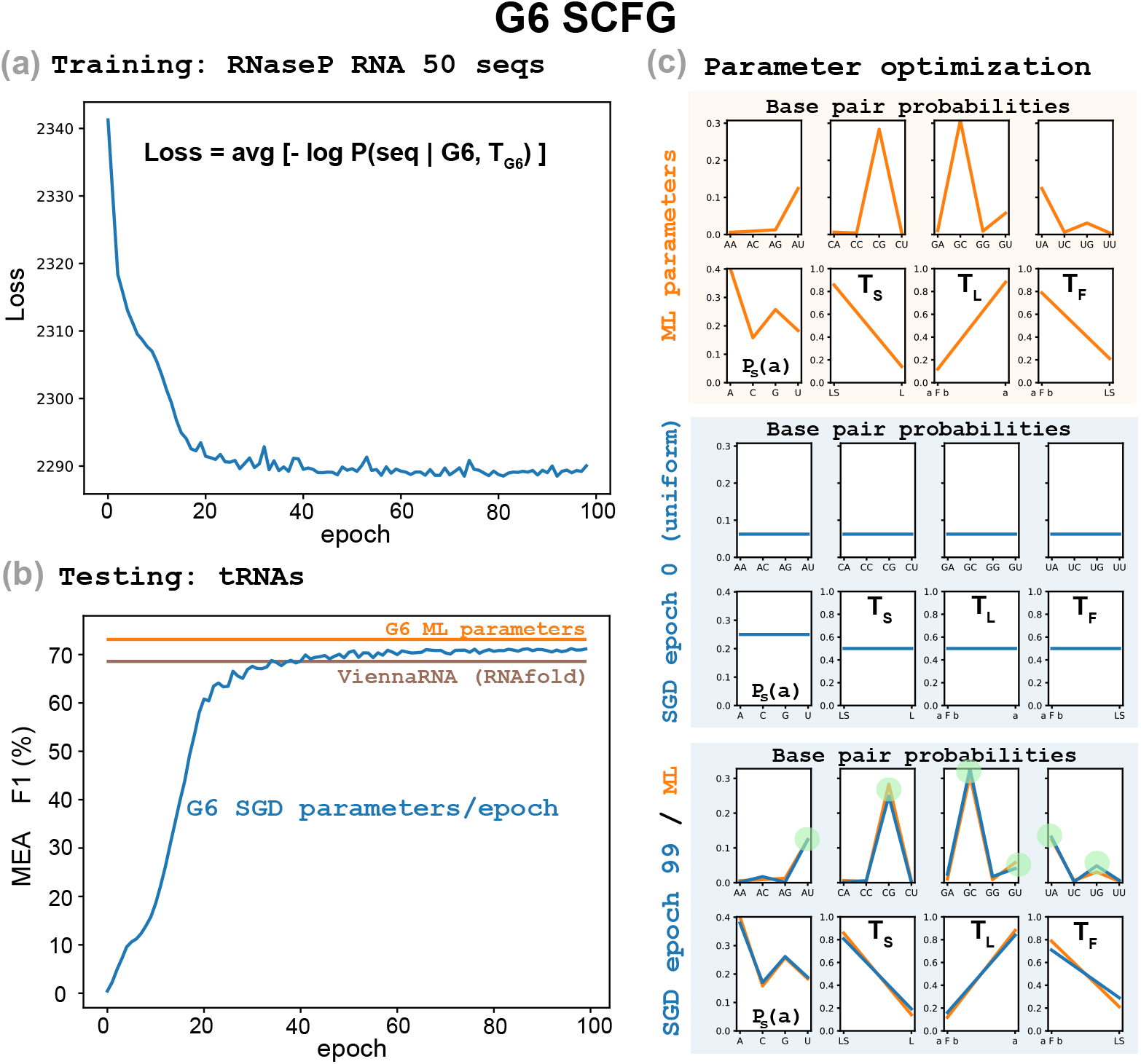
G6 training by SGD on RNA sequences but not structures. The G6 SCFG trained by stochastic gradient descent on a collection of 50 sequences selected at random from a database of RNaseP sequences [6]. **(a)** For a given set of G6 parameters, the loss is defined as the average over the training sequences of the negative log-probability of each sequence (see Eq. 6). **(b)** Performance of the G6 SGD-optimized parameters (per epoch) predicting the secondary structures of a collection of tRNA sequences (blue). We compare to the performance of G6 parameterized by ML which uses both sequence and structure information (orange). We also compare to the predictions using the program RNAfold from ViennaRNA (brown). **(c)** The probability distribution parameters of G6 (base pair probabilities, single residue probability and non-terminal distributions). In orange, parameter values as trained by maximum likelihood (from sequences and structures). In blue, SGD-parameters (trained only from sequences): uniformly distributed at initialization (epoch 0), and after sequence-only SGD optimization (epoch 99, see Eq. 7).

We find that robust learning of base-pairing rules is feasible even with as few as 25–50 sequences, emphasizing how strong the pairwise statistical signal is among structured RNAs.

### Secondary structure prediction is comparable to that of thermodynamic models

To evaluate how well G6 performs in predicting RNA secondary structure when trained only on sequences, we used a random set of 707 tRNAs from the Sprinzl database [27].

To compare structural predictions, we used TORNADO [22]. TORNADO is a general purpose SCFG framework that can accommodate arbitrary SCFGs using customizable parameter sets, and implements most standard structural prediction algorithms. TORNADO allows us to perform a direct comparison between two G6 parameterizations: one trained only on RNaseP RNA sequences, and another trained via maximum likelihood using both RNaseP RNA sequences and structures. In both cases, we report maximum expected accuracy (MEA) predictions.

We measure folding accuracy using the F1 score, which is the harmonic mean of sensitivity (the fraction of true base pairs predicted correctly) and positive predictive value (the fraction of predicted base pairs that are true). F1 is high only if both sensitivity and positive predictive value are high. We report a single global F1 score aggregated across all test sequences.

Figure 2b shows that the G6 grammar trained only on RNaseP sequences performs well on the tRNA dataset (F1 = 71.1% at epoch 99), with accuracy comparable to standard thermodynamic methods such as ViennaRNA (F1 = 68.6%) [14]. It is only slightly worse than the performance of G6 trained by maximum likelihood on both sequences and structures from the RNaseP training set (F1 = 73.1%).

Naturally, performance can get better with much larger models and datasets. For instance, RiNALMo achieves an F1 score of 96.2% on this tRNA test set, compared to our result of F1 = 71.1%. However, the comparison is stark in terms of resources: RiNALMo uses 650 million parameters during pre-training and draws from a large dataset of 102,318 RNA sequences and structures (bpRNA-1m [5]), which includes tRNAs. Even when RiNALMo’s structure head is trained on the smaller ArchiveII dataset [26] excluding all tRNAs (F1 = 94.3%), the tRNA sequences had still been encountered during pre-training. In contrast, our result (F1 = 71.1%) is based on a model with only 21 parameters, trained from scratch on just 50 RNaseP RNA sequences, without access to any structural annotations.

### Base pairing inference does not require aligned sequences

Alignments of RNA sequences are a well-established source of structural information, as the co-evolution found in aligned base paired positions can be exploited to learn quite accurately the base pairs present in a conserved structural RNA [21].

The RNaseP sequences used in our experiment in Figure 2 are not aligned. However, since the training set is made of all homologous RNaseP sequences (with lengths ranging from 189 to 475 nts), there is the possibility that sequences appear quasi-aligned, and that the optimization algorithm learns from some residual covariation.

To challenge the hypothesis of the signal being the result of quasi-aligned homologous RNAs, we expanded the training set to include sequences belonging to seven different structural RNA families. In addition to RNaseP RNAs, the new training set includes: SRP RNAs, tmRNAs, telomerase RNAs, Group I and Group II introns, and 5S rRNA. In order to keep the two training sets to be otherwise comparable in the number of sequences from a given family, we selected a random subset of 400 RNAs from the total 7 Family collection of RNA sequences.

In Figure 3, we observe similar results when using this combined training set. The WCF rules appear as strong and as early as in the RNaseP-alone training case.

**Figure 3:**
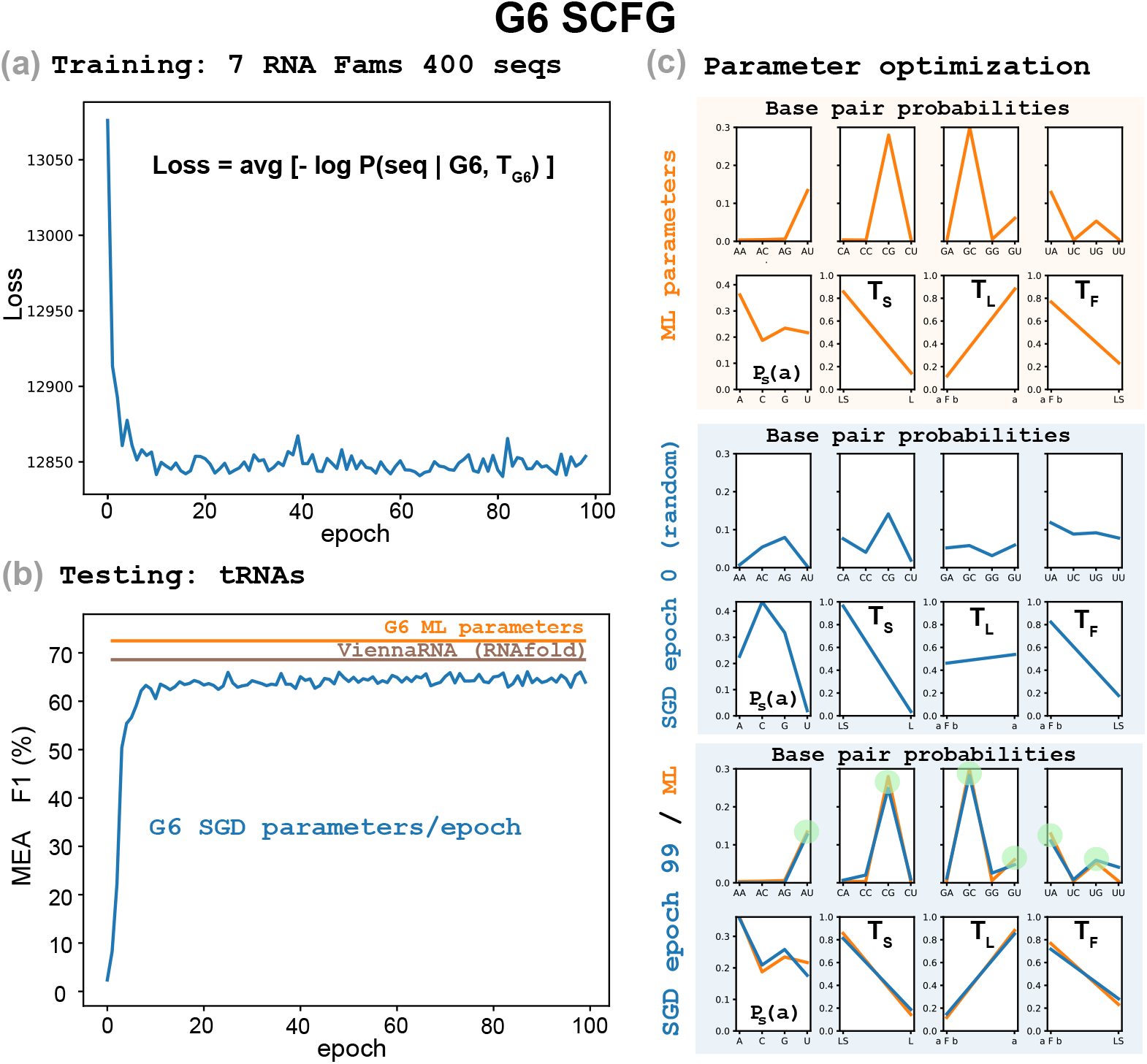
G6 SGD training on non-homologous structural RNA sequences. The G6 SCFG trained by stochastic gradient descent on a sample of 400 sequences selected at random from a collection of 729 sequences from 7 different non-homologous structural RNA families: RNaseP RNA, SRP RNA, tmRNA, telomerase RNA, Group I intron, Group II intron and 5S rRNA. Parameter optimization is performed as described in Eq. 7. **(a)** For a given set of G6 parameters, the loss is the average over the training sequences of the negative log-probability of each sequence (see Eq. 6). **(b)** Performance of the G6 SGD-optimized parameters (per epoch) on predicting the secondary structures of a collection of tRNA sequences (blue). We compare to the performance of G6 parameterized by ML which uses both sequence and structure information (orange). We also compare to the predictions using the program RNAfold from ViennaRNA (brown). **(c)** The probability distribution parameters of G6 (base pair probabilities, single residue probability and non-terminal distributions). In orange, parameter values as trained by maximum likelihood (from sequences and structures). In blue, SGD-parameters (trained only from sequences): uniformly distributed at initialization (epoch 0), and after sequence only SGD-optimization (epoch 99, see Eq. 7). (Results for a uniform distribution initialization, provided in the Supplemental Material, produced similar results.)

### Affine base pair aggregation into helices is key and can be learned just from sequences

Our results show that for structural RNAs, sequence signals for pairwise complementarity are strong and consistent enough across families to enable parameter inference using only sequence-level data, without requiring annotated structures or alignments. Moreover, we wanted to investigate which features in the G6 model are responsible for that result, for which we compared to SCFGs even more reduced than G6.

We selected G6 as one of the smallest SCFGs able to represent RNA secondary structure, and with quite competitive performance compared to other more complex models [6]. There is another SCFG tested for RNA secondary structure, named G5, which has one fewer parameter than G6. The G5 SCFG requires only one non-terminal *S* with rules,

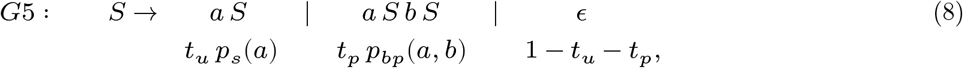

that assign probability 0 ≤ *t*_*p*_ ≤ 1 to a base pair, and 0 ≤ *t*_*u*_ ≤ 1 to an unpaired residue.

It has been shown that trained by maximum likelihood, the G6 grammar performs significantly better than G5 [6]. The key difference that makes G6 work but not G5 is that G6 has two independent probability parameters to control base pairing: *t*_*L*_ controls the start of a helix, and *t*_*F*_ controls adding base pairs to a existing helix (Figure 1), while G5 has only one parameter, *t*_*p*_ (Eq. 8). By selecting parameters that satisfy *t*_*L*_ < *t*_*F*_, G6 is able to give a higher probability to a helix with several base pairs stacked together than to the same number of lone single base pairs. On the other hand, G5 is forced to assigns the same probability to every base pair regardless of whether the base pair occurs as part of a helix or by itself, which is not a realistic description of RNA structure.

Consistently, when we train G5 by SGD, because G5 does not have any incentive to optimize for helices, it is also unable to find the WCF base pair rules, as we show in Figures 4a-4b. We observe that by the time the loss stabilizes, the optimal base pair probabilities do not reproduce WCF base pairing.

**Figure 4:**
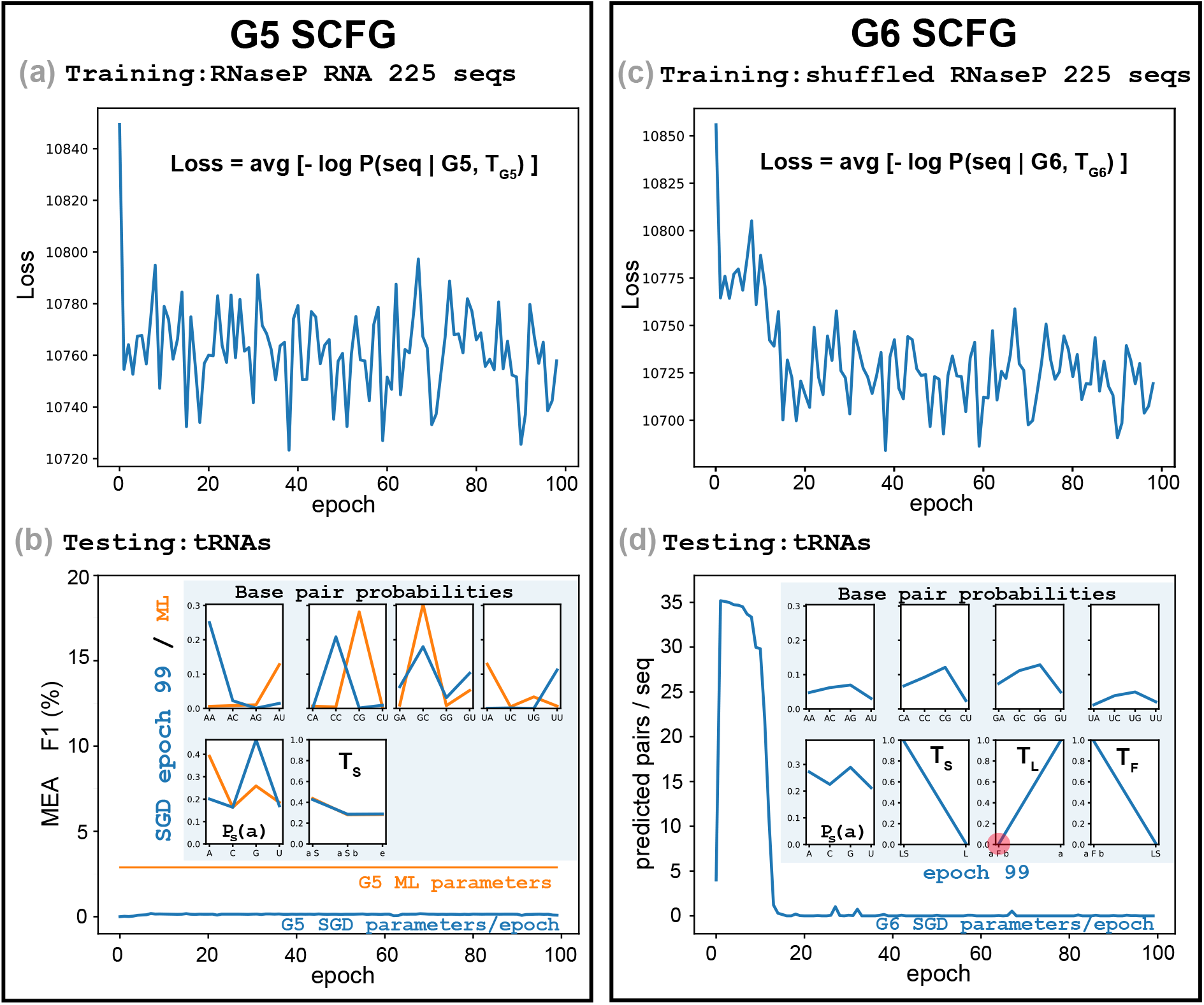
SGD training on a different SCFG (a)-(b), and on random sequences (c)-(d). **(a)** The G5 grammar, with 20 independent parameters, trained on the database of 225 RNaseP RNA sequences. **(b)** Testing on the tRNA dataset with the G5 SGD parameters for each epoch. Insert includes the G5 SGD parameters at epoch 99 (blue), and the maximum likelihood parameters trained from sequences and structures (orange). **(c)** The G6 grammar trained on a shuffled instance of the training set consisting of 225 RNaseP sequences. **(d)** Testing on the tRNA dataset with the G6 SGD parameters for each epoch. The insert includes the optimized probability distribution of G6 at epoch 99. In red, we highlight the value of the rule *L* → *aF b* which initiates a helix (*t*_*L*_ = 0.00418 at epoch 99).

Trained by ML from structures, the G5 base pair probabilities are forced to represent WCF base pairing. However, WCF base pairing does not improve performance for G5. This suggests that it is the combination of WCF base pairing together with the ability to group base pairs into helices what minimally guarantees a model able to learn RNA secondary structure. G6 appears to be the minimal SCFG able to achieve both goals.

It is remarkable that both basic properties of RNA secondary structure can be learned together directly from sequences. Our SGD training procedure that does not see structures, when trained on G6 in addition to learning the WCF base pairs rules, converges to values of the two helix parameters that clearly favor the helical stacking of base pairs that is consistently observed in all RNA structures. In fact, the parameters values of G6 as trained by SGD on the dataset of 50 RNaseP sequences are quite similar to those obtained by ML training on sequences and structures of the same dataset (SGD values: *t*_*L*_ = 0.161, *t*_*F*_ = 0.710; ML values: *t*_*L*_ = 0.119, *t*_*F*_ = 0.788).

Table 1 summarizes the key RNA secondary structure models evaluated in this work, listing model size, training requirements, and performance (F1 score) on the tRNA test set.

**Table 1:**
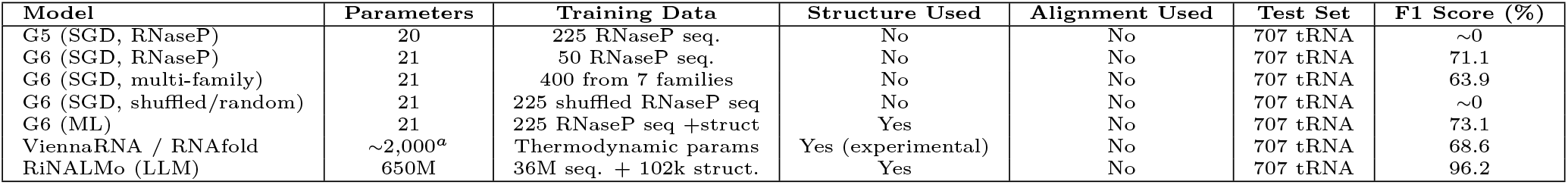
Comparison of RNA secondary structure models and performance on the tRNA test set. ^*a*^Approximate parameter count for ViennaRNA, depends on version. The 7 structural RNA families are: RNaseP RNA, SRP RNA, tmRNA, telomerase RNA, Group I intron, Group II intron and 5S rRNA. F1 scores from main text and Figures; see Methods for training and test sets descriptions.

| Model | Parameters | Training Data | Structure Used | Alignment Used | Test Set | F1 Score (%) |
| --- | --- | --- | --- | --- | --- | --- |
| G5 (SGD, RNaseP) | 20 | 225 RNaseP seq. | No | No | 707 tRNA | ~0 |
| G6 (SGD, RNaseP) | 21 | 50 RNaseP seq. | No | No | 707 tRNA | 71.1 |
| G6 (SGD, multi-family) | 21 | 400 from 7 families | No | No | 707 tRNA | 63.9 |
| G6 (SGD, shuffled/random) | 21 | 225 shuffled RNaseP seq | No | No | 707 tRNA | ~0 |
| G6 (ML) | 21 | 225 RNaseP seq +struct | Yes | No | 707 tRNA | 73.1 |
| ViennaRNA / RNAfold | ~2,000<sup>a</sup> | Thermodynamic params | Yes (experimental) | No | 707 tRNA | 68.6 |
| RiNALMo (LLM) | 650M | 36M seq. + 102k struct. | Yes | No | 707 tRNA | 96.2 |

### What about non-structured or protein-coding sequences? Learning sequence-function relationships beyond RNA structure

To better understand the boundaries of this approach, we trained the G6 model on control data such as random (shuffled) sequences and protein-coding mRNA sequences. Failure to produce WCF base pairing on these control data is as important as success on structured RNA in order to show that the learning is robust in the positive case (structured RNA), but the signal either disappears or changes for random and protein-coding sequences.

### Non-functional shuffled sequences optimize to avoid base pairing

We wanted to test the effect of training on unstructured random sequences. To maintain the base composition intact, relative to the original experiment, we created random sequences by shuffling the RNaseP sequences in the training set. We expect that the model learns from the sequences, thus it should learn that these are not structural but random sequence, but how?

Figure 4c shows that for random sequences, the optimization process converges soon to a regime in which the losses oscillate on a stable range. Interestingly, we observe that the grammar evolves to avoid producing any base pair. The first rule of non-terminal L is responsible for starting a new helix. We observe that the probability of that rule, *t*_*L*_, becomes very small (*t*_*L*_ = 0.00418 after 99 epochs). Consequently, the number of predicted base pairs, given in Figure 4d, shows that after some instability they tend to be almost zero.

In Figure 4d, we also observe that the base pair probabilities do not optimize to WCF values. Moreover, as random sequences make the model optimize to the trivial solution of making almost no base pairs, the values of the base pair probabilities become almost irrelevant as they are rarely used.

### mRNA have sequence-specific pairwise correlations

Finally, we sought to understand what kind of signal the model would capture for a selected set of functional sequences with a signal different from that of structural RNAs, such as messenger RNAs (mRNAs).

We randomly selected two intronless protein coding genes from *S. cerevisiae*, HSP12 and SBH2, and curated datasets of their homologs (see “Methods”). In Figure 5c, we show how when G6 is trained on the HSP12 protein-coding sequences, the model is able to identify correlations between the protein residues [3] that can be observed at the mRNA level [10]. In Figure 5d, we also observe that the optimized parameters are different for SBH2 coding sequences, which stresses the fact that mRNA sequence correlations are not stereotyped as RNA structure is.

**Figure 5:**
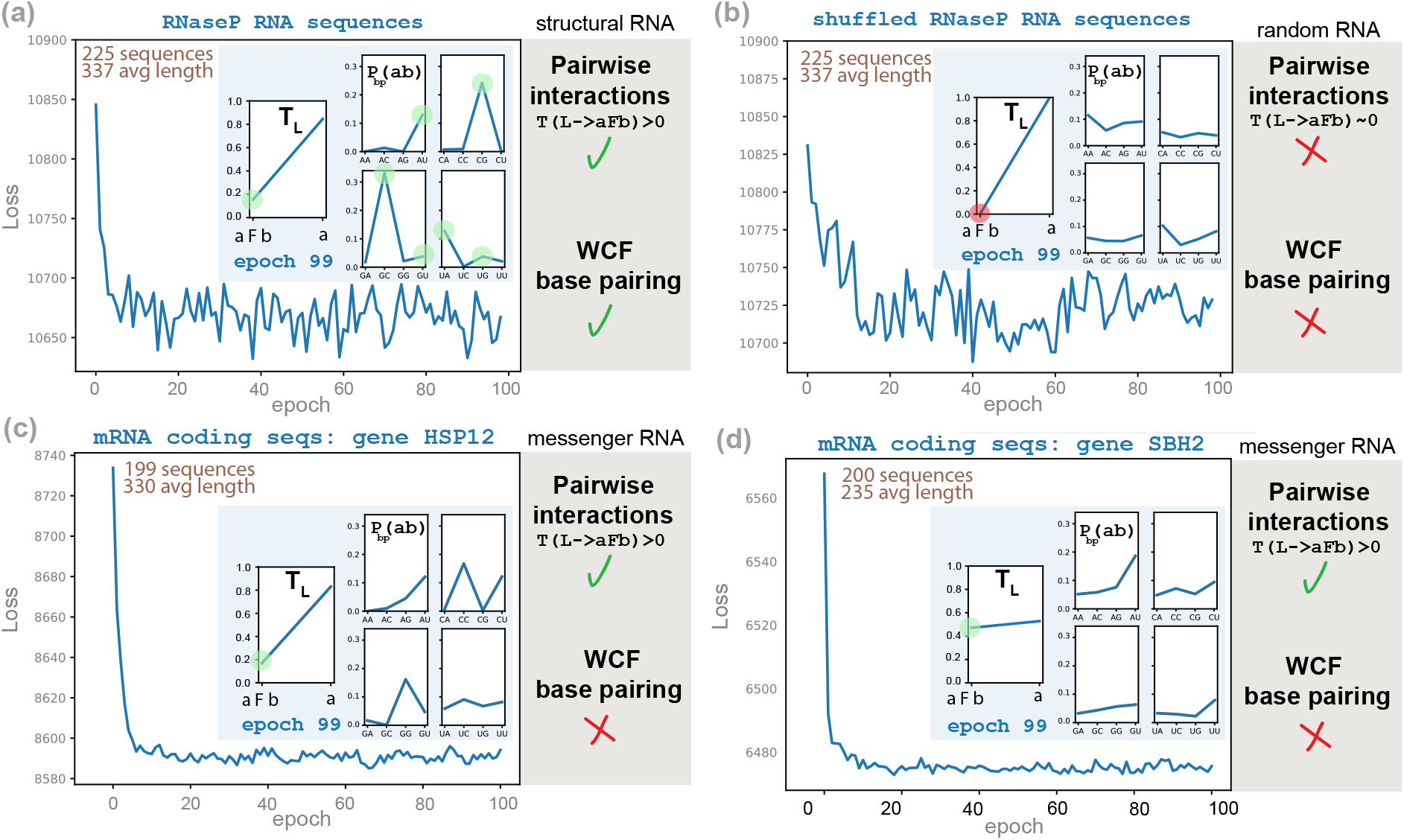
G6 SGD training on sequences with different functions. We compare the results of training the G6 grammar on a similar number of sequences under three different functional constraints: structural RNAs, mRNA sequences, and random sequences. Insert: in blue, parameter values at epoch 99; in green, we highlight when the optimized parameters support the presence of nucleotide interactions, and whether the pairs follow WCF statistics; in red, we indicate when the optimized parameters do not support nucleotide interactions. **(a)** The training set is the database of 225 RNaseP RNAs. **(b)** The training set is a shuffled version of the database of 225 RNaseP RNAs, different from the one in Figures 4c-4d. **(c)** The training set is a collection of 199 homologs of *S. cerevisiae* intronless protein-coding gene HSP12. **(d)** The training set is a collection of 200 homologs of *S. cerevisiae* intronless protein-coding gene SBH2.

There is no equivalent for mRNAs corresponding to the RNA helices of canonical base pairs. When trained on mRNAs, the model captures nonstructural statistical correlations likely reflecting codon usage and amino acid correlations, rather than RNA structural features. This further demonstrates that the emergence of canonical base pairing rules is specific to the organized informational content of structural RNAs, not a universal artifact of SGD optimization.

## Discussion: Implicit functional assumptions

We have introduced a method for training SCFG models for RNA secondary structure via SGD. This method relies on automatic differentiation frameworks for computing the gradients of an objective function with respect to the parameters of an SCFG. Our implementation of SCFGs can automatically be compiled to hardware accelerators such as GPUs.

We have demonstrated that a model with 21 parameters and just the sequences of a few scores of structured RNAs is sufficient to learn the rules of WCF RNA base pairing, as well as the rules of RNA base-pair assembly into helices. Model parameters trained just from sequences produce quite decent secondary structure predictions when compared to standard models with many more parameters such as the thermodynamically parameterized ViennaRNA.

Our results demonstrate that neither structures nor alignments nor models with millions of parameters are necessary to extract the rules of RNA base pairs (Figure 5a). Thus, it should not come as a surprise that large language RNA models not trained on structures or alignments are able to identify base pairs in structural RNAs [4].

On the other hand, what our results also demonstrate is the need to train on structured sequences. In our experiments, we knew ahead of time that RNaseP RNA and the other 6 RNA families are indeed structural RNAs. We built the training sets relying on that information. What kind of results would we have obtained in the absence of that knowledge?

To test that, we trained G6 on sequences that are not structural RNAs. Figure 5b shows that when trained on random sequences, the optimization converges to G6 model parameters that reject the presence of base pairs, which is the result that one would expect and was observed already in Figures 4c-4d for a different random sample. We also trained G6 on two datasets of homologous protein-coding sequences (Figures 5c-5d). The resulting parameters exhibit correlations that are (i) distinct from those inferred from structured RNAs, and (ii) dependent on the target protein. For a given set of RNAs, these results underscore the relationship between conserved structure or function (or lack thereof) and a statistical signature identified as the trained generative model.

Our shallow model trained just on sequences allows us to learn different functional properties, but only after we had separated the sequences by their function. It is now time to expand our simple model in order to be able to infer different sequence functionalities.

### Opportunities for large probabilistic models

The failure to learn WCF base pairs from random or coding sequences confirms that the statistical signal is truly characteristic of structural RNAs. This specificity highlights both the value and the limits of minimal, alignment-free sequence approaches: detection of “structure-like” statistical coupling is robust, but discernment of other functional signals requires at least more expressive grammars, likely beyond the SCFG category, and it may also likely require explicit alignment/covariation information.

While G6 can learn the properties of RNA base pairing, as well as those of particular mRNAs, or even random sequences, obviously it cannot accommodate all of the different interaction patterns observed in functional RNAs at once. Many observed transcripts are still uncharacterized. Distinguishing from sequence alone the functional characteristics of a given transcript or genomic region (even when the answer is that it is just circumstantial transcription without any direct function) is still a work in progress in the field. We hypothesize that probabilistic models with an increased number of parameters trained from sequences alone should be able to classify the functional category of any input biological sequence.

## Methods

### Implementation details

The differentiable JAX implementations of the Inside algorithms for G6 and G5 operate in logarithmic space, and the parameters are normalized after each iteration. We used a learning rate of 0.1. Each iteration used 10 sequence batches selected at random from the training dataset. Explicitly, the grammar parameters *t*_*S*_, *t*_*L*_, *t*_*F*_, *p*_*s*_, *p*_*bp*_ (see Eq. 1) were initialized using the maximum entropy principle to a discrete uniform probability distribution by default. We also used random initializations when specified, with similar results. The code is implemented as part of R-scape v2.6.0, and is provided in the Supplemental Material.

We used TORNADO v0.9.1 [22] to estimate parameters by ML from RNA structures, and to predict secondary structures by maximal expected accuracy using our customized trained parameters. TORNADO is also provided as part of the R-scape v2.6.0 software package.

We used RNAfold (with option `--MEA`) from ViennaRNA v2.7.0 [14], and RiNALMo [17] with the largest model available (“giga”, 650 million parameters pre-trained on 36 million non-coding RNA sequences) on a NVIDIA A100-SXM4-40GB through Google Colab. The RiNALMo structural module was trained on bpRNA-1m with 102,318 RNA structures [5], and on the ArchiveII dataset with 3,975 structures (https://rna.urmc.rochester.edu/pub/archiveII.tar.gz) [26], after all tRNAs had been removed.

### Datasets

The RNA training datasets include: a collection of 225 RNaseP RNA sequences (avg. length 337 nts) from Dowell&Eddy [6], selected from the Ribonuclease P database [2]. Sequences from six other structural RNA families: 81 SRP RNAs, 97 tmRNAs, 37 telomerase RNAs, 16 Group I introns, 3 Group II introns, and 279 5S rRNAs, with annotated secondary structures previously collected by TORNADO from Dowell&Eddy [6] and the Archive database (https://rna.urmc.rochester.edu/pub/archiveII.tar.gz).

For the training set in Figure 2, we used a subsample of 50 RNaseP RNA sequences from the RNaseP dataset above. For the training set in Figure 3, 400 sequences with lengths up to 400 nts were selected at random from the total set of 729 sequences from all 7 RNA families. The test dataset is a collection of 707 tRNA sequences (avg. length 74 nts) from the Sprinzl *et al*. dataset [27].

For the mRNA training datasets in Figures 5c-5d, we used *S. cerevisiae* (strain S288C) intronless protein coding genes: HSP12 (chrVI, YFL014W) and SBH2 (chrV, YER019C-A). The genes were selected at random from the set of 304 intronless genes in *S. cerevisiae* that are shorter than 400 nts. We used nhmmer [28], to identify homologs in other *Ascomycota* fungal genomes, from which we selected a random subset of up to 200 homologs per mRNA (out of 199 for HSP12 and 248 for SBH2).

All datasets are provided in the Supplemental Material.

## Acknowledgments

We thank Marcell Szikszai for help running the software RiNALMo, and Max Ward for insights into automatic differentiation of RNA folding models. We thank William Gao for providing the fungal mRNA sequences. We thank Sean R. Eddy and William Gao for a critical reading of the manuscript. E. R. acknowledges the hospitality of the Centro de Ciencias de Benasque Pedro Pascual, Benasque, Spain, during the completion of this manuscript.

## Author Contributions

E.R. conceived the research. J.S.P. and R.K.K. implemented the algorithms for the G5 grammar. E.R. implemented the algorithms for the G6 grammar. E.R. performed the experiments and wrote the manuscript. All authors edited the manuscript.

## Extended Data Figures

**Figure S1:**
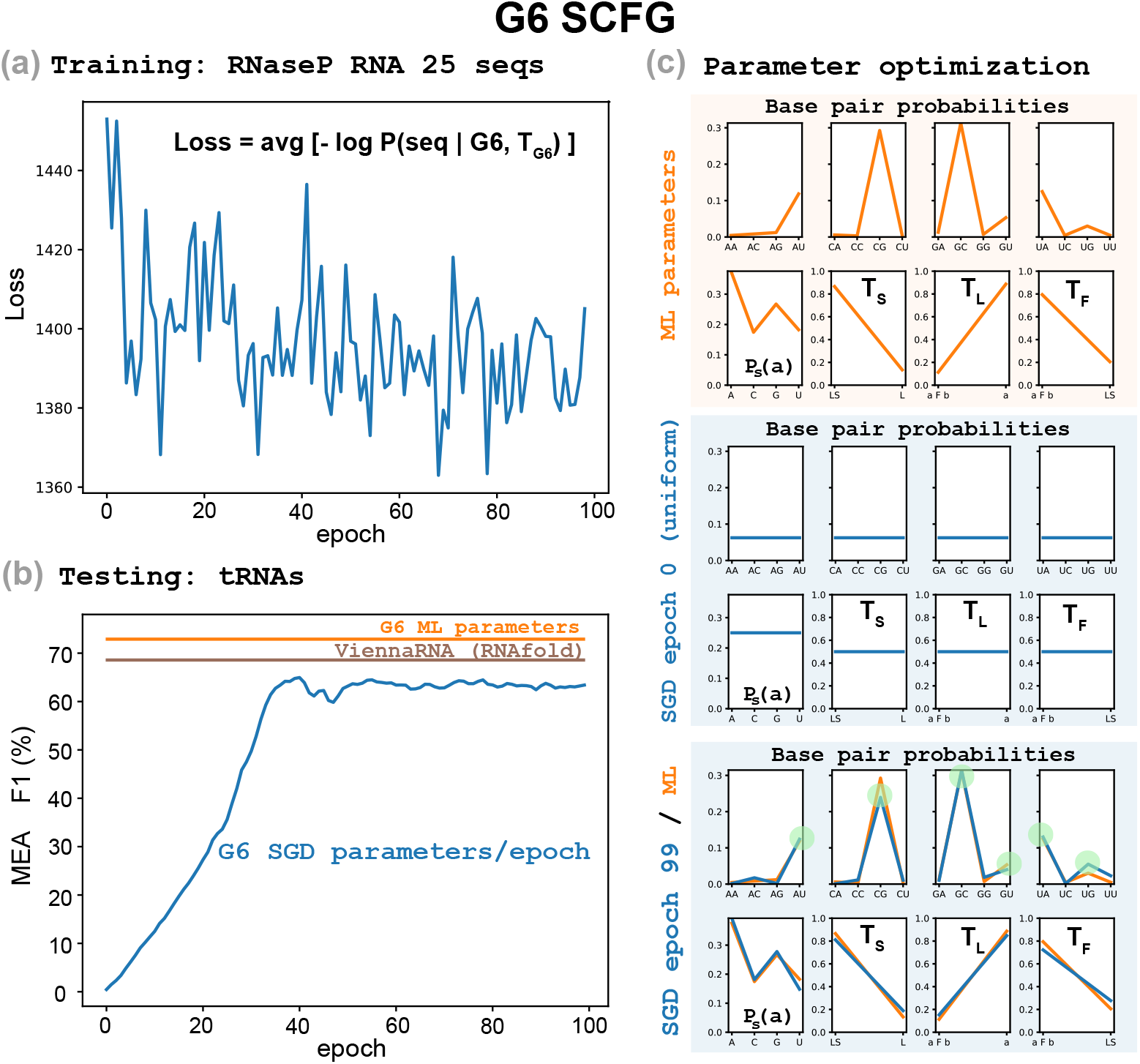
G6 SGD training on 25 RNaseP RNA sequences. We tested performance of our SGD optimization algorithm on a random subsample of 25 RNaseP RNA sequences. Legends for the different sections are similar to those in Figure 2.

