## Supplementary material for "What does it take to learn the rules of RNA base pairing? A lot less than you may think": coding_all_mRNA_gene_info.pl

```
#!/usr/bin/perl -w
# 
#  Rank a R-scape output by the F measure
#
use strict;
use Class::Struct;
use POSIX;

use List::Util qw/shuffle/;

use constant GNUPLOT => '/usr/local/bin/gnuplot';
use lib '/Users/erivas/src/Mysrc/R-scape/scripts';
use FUNCS;

use vars qw ($opt_b $opt_p $opt_v $opt_f $opt_F);  # required if strict used
use Getopt::Std;
getopts ('b:p:vf:F:');

# Print a helpful message if the user provides no input file.
if (!@ARGV) {
    print "usage:  coding.pl [options]  <chr> <max_len> \n\n";
    print "options:\n";
    print "-v    :  be verbose\n";
    exit;
}
my $chr     = shift;
my $max_len = shift;

my $mRNA_file = "/Users/erivas/projects/d-SCFG_June2025/data/coding/all_mRNA_gene_info.txt";
my $fafile   = ($max_len > 0)?
    "/Users/erivas/projects/d-SCFG_June2025/data/coding/$chr.len$max_len.fa" :
    "/Users/erivas/projects/d-SCFG_June2025/data/coding/$chr.fa";
print ("file    $mRNA_file\n");
print ("outfile $fafile\n");
open (OUT, ">$fafile") || die;

my $this_chr;
my $this_name;
my $this_seq;
my $nseq = 0;

open(FILE, "$mRNA_file") || die;
while (<FILE>) {
    if (/\#/) {
    }
    elsif (/^(chr\S+)\s+(\S+)\s+\d+\s+\d+\s+(\S+)$/) {
	$this_chr  = $1;
	$this_name = $2;
	$this_seq  = ($max_len > 0)? substr($3, 0, $max_len) : $3;
	
	if ($chr =~ /^$this_chr$/) {
	    $nseq ++;
	    print ">$this_name\n$this_seq\n";
	    print OUT ">$this_name\n$this_seq\n";
	}
    }
}
close(FILE);
print("chr $chr nsq $nseq\n");

close(OUT);

my $sto_file = fasta_to_sto($fafile);

sub fasta_to_sto{
    my ($fastafile) = @_;

    my $filename = $fastafile;
    if ($filename =~ /^(\S+)\.fa/) { $filename = $1; }
    
    my $stofile = "$filename.sto";
    open (OUT, ">$stofile") || die;
    
    my $nsq = 0;
    my @sq;
    my @sqname;
    FUNCS::parse_afafile($fastafile, \$nsq, \@sq, \@sqname);
    for (my $s = 0; $s < $nsq; $s ++) {
	printf OUT "# STOCKHOLM 1.0\n\n"; 
	printf OUT "%s %s\n", $sqname[$s], $sq[$s]; 
	printf OUT "//\n"; 
    }
    
    close(OUT);
    return $stofile;
    
}
```
