## Supplementary material for "What does it take to learn the rules of RNA base pairing? A lot less than you may think": conus_select.pl

```
#!/usr/bin/perl -w
# 
#  Rank a R-scape output by the F measure
#
use strict;
use Class::Struct;
use POSIX;

use List::Util qw/shuffle/;

use constant GNUPLOT => '/usr/local/bin/gnuplot';
use lib '/Users/erivas/src/Mysrc/R-scape/scripts';
use FUNCS;

use vars qw ($opt_b $opt_p $opt_v $opt_f $opt_F);  # required if strict used
use Getopt::Std;
getopts ('b:p:vf:F:');

# Print a helpful message if the user provides no input file.
if (!@ARGV) {
    print "usage:  conus_select.pl [options] <n> <stofile> \n\n";
    print "options:\n";
    print "-v    :  be verbose\n";
    exit;
}
my $n         = shift;
my $extra     = shift;
my $conus_sto = shift;

my $name;
my $dir;
if ($conus_sto =~ /^(\S+)\/([^\/]+)\.sto/) { $dir = $1; $name = $2; }
else { print "bad file $conus_sto\n"; die; }
print "dir:$dir\n";
print "name:$name\n";


my $esl_reformat = "~/src/Mysrc/R-scape/lib/hmmer/easel/miniapps/easel reformat";

my $sto_select_n      = sto_select_random($n, $extra, $dir, $name, $conus_sto);
my $sto_select_n_noss = sto_to_noss($sto_select_n);
my $fasta_select_n    = sto_to_fasta($sto_select_n);


sub sto_select_random{
    my ($n, $extra, $dir, $name, $stofile) = @_;
    
    my $select_sto_file = "$dir/$name.$n.$extra.sto";
    print "new sto: $select_sto_file\n";
    open (OUT, ">$select_sto_file") || die;
    
    my $nsq = 0;
    my @sq;
    my @ss;
    my @sqname;
    
    my $thisname;
    my $thissq;
    my $thisss;
    open (STO, "$stofile") || die;
    while (<STO>) {
	if (/# STOCKHOLM 1.0/) {
	    $thissq = "";
	    $thisss = "";
	}
	elsif (/^([^\#]\S+)\s+(\S+)$/) {
	    $thisname  = $1;
	    $thissq   .= $2;
	}
	elsif (/#=GR\s+$thisname\s+SS\s+(\S+)$/) {
	    $thisss  .= $1;
	}
	elsif (/\/\//) {
	    printf "# STOCKHOLM 1.0\n\n"; 
	    printf "%s         %s\n", $thisname, $thissq; 
	    printf "#=GR %s SS %s\n", $thisname, $thisss; 
	    printf "//\n"; 

	    $sqname[$nsq] = $thisname;
	    $sq[$nsq]     = $thissq;
	    $ss[$nsq]     = $thisss;
	    $nsq ++;
	}
    }
    close(STO);
    
    if ($n > $nsq) { $n = $nsq; }
    
    my @shuffled_idx = shuffle(0..$nsq-1);
    my @pick_idx     = @shuffled_idx[0 .. $n-1];
    
    my @pick_sq     = @sq    [ @pick_idx ];
    my @pick_ss     = @ss    [ @pick_idx ];
    my @pick_sqname = @sqname[ @pick_idx ];
    
  
    for (my $s = 0; $s < $n; $s ++) {
	printf OUT "# STOCKHOLM 1.0\n\n"; 
	printf OUT "%s         %s\n", $pick_sqname[$s], $pick_sq[$s]; 
	printf OUT "#=GR %s SS %s\n", $pick_sqname[$s], $pick_ss[$s]; 
	printf OUT "//\n"; 
     }
    close(OUT);
    
    return $select_sto_file;
}


sub fasta_select_random{
    my ($n, $dir, $name, $file) = @_;
    
    my $select_fasta_file = "$dir/$name.$n.fa";
    open (OUT, ">$select_fasta_file") || die;
    
    my $nsq = 0;
    my @sq;
    my @sqname;
    FUNCS::parse_afafile($file, \$nsq, \@sq, \@sqname);

    if ($n > $nsq) { $n = $nsq; }
    
    my @shuffled_idx = shuffle(0..$nsq-1);
    my @pick_idx     = @shuffled_idx[0 .. $n-1];
    
    my @pick_sq     = @sq    [ @pick_idx ];
    my @pick_sqname = @sqname[ @pick_idx ];
    
    
    for (my $s = 0; $s < $n; $s ++) {
        printf OUT ">%s\n%s\n", $pick_sqname[$s], $pick_sq[$s]; 
    }
    close(OUT);
    
    return $select_fasta_file;
}


sub fasta_to_sto{
    my ($fastafile) = @_;

    my $filename = $fastafile;
    if ($filename =~ /^(\S+)\.fa/) { $filename = $1; }
    
    my $stofile = "$filename.sto";
    open (OUT, ">$stofile") || die;
    
    my $nsq = 0;
    my @sq;
    my @sqname;
    FUNCS::parse_afafile($fastafile, \$nsq, \@sq, \@sqname);
    for (my $s = 0; $s < $nsq; $s ++) {
	printf OUT "# STOCKHOLM 1.0\n\n"; 
	printf OUT "%s %s\n", $sqname[$s], $sq[$s]; 
	printf OUT "//\n"; 
    }
    
    close(OUT);
    return $stofile;
}


sub sto_to_fasta{
    my ($stofile) = @_;

    my $filename = $stofile;
    if ($filename =~ /^(\S+)\.sto/) { $filename = $1; }
    
    my $fasta_file = "$filename.fa";

    my $cmd;
    $cmd = "$esl_reformat fasta $stofile > $fasta_file";
    system("$cmd \n");

    return $fasta_file;
}

sub sto_to_noss{
    my ($stofile) = @_;

    my $filename = $stofile;
    if ($filename =~ /^(\S+)\.sto/) { $filename = $1; }
    
    my $noss_file = "$filename.noss.sto";

    my $cmd;
    $cmd = "cat $stofile | grep -v SS > $noss_file";
    system("$cmd \n");

    return $noss_file;
}
```
