## Supplementary material for "What does it take to learn the rules of RNA base pairing? A lot less than you may think": RINALMo.pl

```
#!/usr/bin/perl -w
# 
#  Rank a R-scape output by the F measure
#
use strict;
use Class::Struct;
use POSIX;
    
use constant GNUPLOT => '/usr/local/bin/gnuplot';
use lib '/Users/erivas/src/Mysrc/R-scape/scripts';
use FUNCS;

use vars qw ($opt_b $opt_p $opt_v $opt_f $opt_F);  # required if strict used
use Getopt::Std;
getopts ('b:p:vf:F:');

# Print a helpful message if the user provides no input file.
if (!@ARGV) {
    print "usage:  RINALMo.pl [options] <true.sto> <RINALMo.out> \n\n";
    print "options:\n";
    print "-v    :  be verbose\n";
    exit;
}
my $stofile     = shift;
my $RINALMo_dir = shift;

my $RINALMo_sto = RINALMo_ct_to_sto($RINALMo_dir, $stofile);

my $esl_compstruct = "~/src/Mysrc/R-scape/lib/hmmer/easel/miniapps/esl-compstruct";

my $RINALMo_acc = "$RINALMo_sto.acc";

my $cmd;
$cmd = "$esl_compstruct $stofile $RINALMo_sto > $RINALMo_acc";
system("$cmd \n");

parse_acc($RINALMo_acc);


sub parse_acc {
    my ($acc_file) = @_;
    
    my $F   = 0;
    my $sen = 0;
    my $ppv = 0;
    
    #Overall prediction accuracy (1415 sequences, 104851 positions)
    #29469/29469 trusted pairs predicted (100.00% sensitivity)
    #29469/29469 predicted pairs correct (100.00% PPV)
    
    open(FILE, "$acc_file") || die;
    while(<FILE>) {
        if (/^Overal prediction/) {
        }
	elsif (/trusted pairs predicted \((\d+\.\d+)\% sensitivity/) {
	    $sen = $1;
	}
	elsif (/predicted pairs correct \((\d+\.\d+)\% PPV/) {
	    $ppv = $1;
	}
    }
    close(FILE);
    
    if ($sen + $ppv > 0) { $F = 2*$sen*$ppv / ($sen + $ppv); }
    
    
    print("$acc_file\n");
    print("F $F sen $sen ppv $ppv\n");
}


sub RINALMo_ct_to_sto{
    
    my ($RINALMo_dir, $stofile) = @_;
    
    my $RINALMo_name = "$RINALMo_dir";
    if ($RINALMo_name =~ /\/([^\/]+)\s*$/) { $RINALMo_name = $1; }
    
    my $RINALMo_sto = "$RINALMo_dir/$RINALMo_name.sto";
    print("RINALMo: $RINALMo_sto\n");
    open(STO, ">$RINALMo_sto") || die;
    
    my $nct = 0;
    my @name;
    my @seq;
    my @ss;
    
    opendir my $dir, $RINALMo_dir or die "Cannot open directory: $!";
    my @ct_files = readdir $dir;
    
    foreach my $ct_file (@ct_files) {
	if ($ct_file =~ /^(\S+)\.ct/) {
	    my $name = "$1";
	    my $seq  = "";
	    my $ss   = "";
	    $ct_file = "$RINALMo_dir/$ct_file";
	    print("$nct $ct_file\n");
	    
	    parse_ct("$ct_file", \$seq, \$ss);
	    print "$seq\n$ss\n";
	    
	    $name[$nct] = $name;
	    $seq[$nct]  = $seq;
	    $ss[$nct]   = $ss;
	    $nct ++;
	}
    }
    closedir $dir;
    
    # parse sto file for name order
    my $n_sto = 0;
    my @name_input;

    my $n_lines = 0;
    open(FILE, "$stofile") || die;
    while(<FILE>) {
	if (/\#=GR\s+(\S+)\s+SS/) {
	    if ($n_lines == 0) {
		$name_input[$n_sto] = $1;
		$n_sto ++;
		$n_lines ++;
	    }
	}
	elsif (/\/\//) {
	    $n_lines = 0;
	}
    }
    close(FILE);
    
    if ($nct != $n_sto) { print(" nct $nct should be n_sto $n_sto\n"); die;}
    
    
    for (my $n = 0; $n < $n_sto; $n ++) {
	my $name_i = $name_input[$n];
	
	my $m;
	for ($m = 0; $m < $n_sto; $m ++) {
	    
	    if ($name_i =~ /^($name[$m])$/) {
		print STO "# STOCKHOLM 1.0\n\n";
		print STO "$name_i         $seq[$m]\n";
		print STO "#=GR $name_i SS $ss[$m]\n";
		print STO "\/\/\n";
		
		last;
	    }
	}
	if ($m == $n_sto) { print("$name_i not found\n"); die; }	
    }
    
    close(STO);
    return $RINALMo_sto;
}


sub parse_ct {
    my ($ct_file, $seq_ret, $ss_ret) = @_;
    
    my $seq = "";
    my $ss  = "";
    
    my $len;
    my @ct;
    my $n = 0;
    my $which;
    open(FILE, "$ct_file") || die;
    while(<FILE>) {
	if (/^(\d+)\s+(\S+)\s+\d+\s+\d+\s+(\d+)\s+\d+\s*$/) {
	    $which      = $1 - 1;
	    $seq       .= $2;
	    $ct[$which] = $3 - 1;
	    $n ++;
	}
        elsif (/^(\d+)\s+\S+/) {
	    $len = $1;
	}
    }
    close(FILE);

    FUNCS::ct2ss($len, \@ct, \$ss);
    #print "$seq\n$ss\n";
    
    $$seq_ret = $seq;
    $$ss_ret  = $ss;
    
}
```
