## Supplementary material for "What does it take to learn the rules of RNA base pairing? A lot less than you may think": S288C_intronless_select.pl

```
#!/usr/bin/perl -w
# 
#  Rank a R-scape output by the F measure
#
use strict;
use Class::Struct;
use POSIX;

use List::Util qw/shuffle/;

use constant GNUPLOT => '/usr/local/bin/gnuplot';
use lib '/n/eddy_lab/users/erivas/rscape_v2.5.8/scripts';
use FUNCS;

use vars qw ($opt_b $opt_p $opt_v $opt_f $opt_F);  # required if strict used
use Getopt::Std;
getopts ('b:p:vf:F:');

# Print a helpful message if the user provides no input file.
if (!@ARGV) {
    print "usage:  S288C_intronless_select.pl [options] <maxlen>  \n\n";
    print "options:\n";
    print "-v    :  be verbose\n";
    exit;
}

my $maxlen = shift;

my $wgaodir = "/n/eddy_lab/users/wgao/S288C_data/mRNA_related/intronless_genes";
my $outdir  = "~/projects/d-SCFG_June2025/data/coding";

my $ntotal = 0;

$ntotal += parse_dir("chrI");
$ntotal += parse_dir("chrII");
$ntotal += parse_dir("chrIII");
$ntotal += parse_dir("chrIV");
$ntotal += parse_dir("chrV");
$ntotal += parse_dir("chrVI");
$ntotal += parse_dir("chrVII");
$ntotal += parse_dir("chrVIII");
$ntotal += parse_dir("chrIX");
$ntotal += parse_dir("chrX");
$ntotal += parse_dir("chrXI");
$ntotal += parse_dir("chrXII");
$ntotal += parse_dir("chrXIII");
$ntotal += parse_dir("chrXIV");
$ntotal += parse_dir("chrXV");
$ntotal += parse_dir("chrXVI");
    
print ("ngenes len < $maxlen total = $ntotal\n");

sub parse_dir {
    my ($chr) = @_;
	
    my $chrdir = "$wgaodir/$chr";

    my $n_genes = 0;
    opendir my $dir, $chrdir or die "Cannot open directory: $!";
    my @files = readdir $dir;
    
    foreach my $file (@files) {
	
	if ($file =~ /^(\S+)\.fasta/) {
	    my $name = $1;
	    my $fastafile = "$chrdir/$file";
	    
	    my $nsq = 0;
	    my @sq;
	    my @sqname;
	    FUNCS::parse_afafile($fastafile, \$nsq, \@sq, \@sqname);
	    if ($nsq != 1) { print("wrong file?\n"); die; }
	    
	    my $len = length($sq[0]);
	    
	    if ($len <= $maxlen) {
		my $stofile = "$chrdir/$name/rm_v1.sto";
		my $sto_nsq;
		my @sto_name;
		my @sto_sq;
		my $sto_ss;
		my @sto_ct;
		my $sto_rfasq;
		
		FUNCS::parse_stofile($stofile, \$sto_nsq, \@sto_name, \@sto_sq, \$sto_ss, \@sto_ct, \$sto_rfasq, 1);
		print "$chr $name len=$len nseq = $sto_nsq\n";
		$ngenes ++;
	    }
	    
	}
    }
    closedir $dir;
    print ("ngenes len < $maxlen for $chr = $ngenes\n");
    return $ngenes;
}
```
