## Supplementary material for "What does it take to learn the rules of RNA base pairing? A lot less than you may think": tornado.pl

```
#!/usr/bin/perl -w
# 
#  Rank a R-scape output by the F measure
#
use strict;
use Class::Struct;
use POSIX;

use List::Util qw/shuffle/;

use constant GNUPLOT => '/usr/local/bin/gnuplot';
use lib '/Users/erivas/src/Mysrc/R-scape/scripts';
use FUNCS;

use vars qw ($opt_b $opt_p $opt_v $opt_f $opt_F);  # required if strict used
use Getopt::Std;
getopts ('b:p:vf:F:');

# Print a helpful message if the user provides no input file.
if (!@ARGV) {
    print "usage:  tornado.pl [options] <n> <min_len> <max_len> <stofile> \n\n";
    print "options:\n";
    print "-v    :  be verbose\n";
    exit;
}
my $n_shuffle = shift;
my $min_len   = shift;
my $max_len   = shift;
my $sto_file  = shift;

my $name;
my $dir;
if ($sto_file =~ /^(\S+\/)([^\/]+)\.sto/) { $dir = $1; $name = $2; }
else { print "bad file $sto_file\n"; die; }

my $esl_reformat = "~/src/Mysrc/R-scape/lib/hmmer/easel/miniapps/esl-reformat";


# select n from sto -> convert to fasta and noss.sto
# 
my $sto_select_n      = sto_select_n($n_shuffle, $max_len, $dir, $name, $sto_file);
my $sto_select_n_noss = sto_to_noss($sto_select_n);
my $fasta_select_n    = sto_to_fasta($sto_select_n);


sub sto_select_n{
    my ($n, $max_len, $dir, $name, $stofile) = @_;
    
    my $select_sto_file = "$dir/$name.len$min_len\_$max_len.nsq$n.sto";
    open (OUT, ">$select_sto_file") || die;
    
    my $nsq = 0;
    my $nsq_total = 0;
    my @sq;
    my @ss;
    my @name;

    my $this_sq;
    my $this_ss;
    my $this_name;

    print("sto $stofile\n");
    open (STO, "$stofile") || die;
    while (<STO>) {
	if (/\# STOCKHOLM 1.0/) {
	    $this_sq = "";
	    $this_ss = "";
	    $this_name = "";
	}
 	elsif (/\#=GR\s+$this_name\s+SS\s+(\S+)\s*$/) {
	 $this_ss .= $1; 
	}
	elsif (/^(\S+)\s+(\S+)\s*$/) {
	    $this_name  = $1; 
	    $this_sq   .= $2; 	  
	}
	elsif (/\/\//) {
	    $nsq_total ++;
	    # select sequences up to max_len
	    if ($min_len == 0 || length($this_sq) >= $min_len) {
		if ($max_len == 0 || length($this_sq) <= $max_len) {
		    $sq[$nsq]   = $this_sq;
		    $ss[$nsq]   = $this_ss;
		    $name[$nsq] = $this_name;
		    $nsq ++;
		    #print("\n^^$nsq $this_name\n$this_sq\n$this_ss\n");
		}
	    }
	}
    }
    close (STO);
    print("nsq $nsq/$nsq_total min_len $min_len max_len $max_len\n");
    
    if ($n > $nsq) { $n = $nsq; }
    
    my @shuffled_idx = shuffle(0..$nsq-1);
    my @pick_idx     = @shuffled_idx[0 .. $n-1];
    
    my @pick_sq     = @sq  [ @pick_idx ];
    my @pick_ss     = @ss  [ @pick_idx ];
    my @pick_sqname = @name[ @pick_idx ];
    
    for (my $s = 0; $s < $n; $s ++) {
	printf OUT "# STOCKHOLM 1.0\n\n"; 
	printf OUT "%s         %s\n", $pick_sqname[$s], $pick_sq[$s]; 
	printf OUT "#=GR %s SS %s\n", $pick_sqname[$s], $pick_ss[$s]; 
	printf OUT "//\n"; 
    }

    close(OUT);
    
    return $select_sto_file;
}


sub sto_to_fasta{
    my ($stofile) = @_;

    my $filename = $stofile;
    if ($filename =~ /^(\S+)\.sto/) { $filename = $1; }
    
    my $fasta_file = "$filename.fa";

    my $cmd;
    $cmd = "$esl_reformat fasta $stofile > $fasta_file";
    system("$cmd \n");

    return $fasta_file;
}

sub sto_to_noss{
    my ($stofile) = @_;

    my $filename = $stofile;
    if ($filename =~ /^(\S+)\.sto/) { $filename = $1; }
    
    my $noss_file = "$filename.noss.sto";

    my $cmd;
    $cmd = "cat $stofile | grep -v SS > $noss_file";
    system("$cmd \n");

    return $noss_file;
}
```
