## Supplementary material for "What does it take to learn the rules of RNA base pairing? A lot less than you may think": ViennaRNA.pl

```
#!/usr/bin/perl -w
# 
#  Rank a R-scape output by the F measure
#
use strict;
use Class::Struct;
use POSIX;
    
use constant GNUPLOT => '/usr/local/bin/gnuplot';
use lib '/Users/erivas/src/Mysrc/R-scape/scripts';
use FUNCS;

use vars qw ($opt_b $opt_p $opt_v $opt_f $opt_F);  # required if strict used
use Getopt::Std;
getopts ('b:p:vf:F:');

# Print a helpful message if the user provides no input file.
if (!@ARGV) {
    print "usage:  ViennaRNA.pl [options] <true.sto> <vienna.out. \n\n";
    print "options:\n";
    print "-v    :  be verbose\n";
    exit;
}
my $stofile    = shift;
my $viennafile = shift;

my $vienna_mfe_sto = ViennaRNA_to_sto($viennafile, 1);
my $vienna_mea_sto = ViennaRNA_to_sto($viennafile, 0);

my $esl_compstruct = "~/src/Mysrc/R-scape/lib/hmmer/easel/miniapps/esl-compstruct";

my $vienna_mfe_acc = "$vienna_mfe_sto.acc";
my $vienna_mea_acc = "$vienna_mea_sto.acc";

my $cmd;
$cmd = "$esl_compstruct $stofile $vienna_mfe_sto > $vienna_mfe_acc";
system("$cmd \n");
$cmd = "$esl_compstruct $stofile $vienna_mea_sto > $vienna_mea_acc";
system("$cmd \n");

parse_acc($vienna_mfe_acc);
parse_acc($vienna_mea_acc);


sub parse_acc {
    my ($acc_file) = @_;

    my $F   = 0;
    my $sen = 0;
    my $ppv = 0;

   #Overall prediction accuracy (1415 sequences, 104851 positions)
   #29469/29469 trusted pairs predicted (100.00% sensitivity)
   #29469/29469 predicted pairs correct (100.00% PPV)

    open(FILE, "$acc_file") || die;
    while(<FILE>) {
        if (/^Overal prediction/) {
        }
	elsif (/trusted pairs predicted \((\d+\.\d+)\% sensitivity/) {
	    $sen = $1;
	}
	elsif (/predicted pairs correct \((\d+\.\d+)\% PPV/) {
	    $ppv = $1;
	}
    }
    close(FILE);

    if ($sen + $ppv > 0) { $F = 2*$sen*$ppv / ($sen + $ppv); }
    
    
    print("$acc_file\n");
    print("F $F sen $sen ppv $ppv\n");
}


sub ViennaRNA_to_sto{
    my ($viennafile, $mfe) = @_;

    my $viennasto = ($mfe)? "$viennafile.mfe.sto" : "$viennafile.mea.sto";

    my $name     = "";
    my $extra    = "";
    my $fullname = "";
    my $seq  = "";
    my $ss   = "";

    open(STO, ">$viennasto") || die;

    my $nsq = 0;
    open(FILE, "$viennafile") || die;
    while(<FILE>) {
        if (/^>(\S+)(\s.*)$/) {
	    if ($nsq > 0) {
		if ($seq =~ // || $ss =~ //) { print("seq or ss is bad\n"); die; }
		print STO "# STOCKHOLM 1.0\n\n";
		if ($extra =~ /\S+/) {
		    print STO "#=GS $name$extra\n\n";
		}
		print STO "$name         $seq\n";
		print STO "#=GR $name SS $ss\n";
		print STO "\/\/\n";
	    }
	    
	    $name  = $1;
	    $extra = $2;
	    $seq = "";
	    $ss = "";
	    $nsq ++;
        }
	elsif (/^([A-Za-z]\S+)/) {
	    $seq = $1;
	}
	elsif ( $mfe && /^([\(\.]\S+)\s+\(\s*-\d+\.\d+\)/) {
	    $ss = $1;
	}
	elsif (!$mfe && /^([\(\.]\S+)\s+\{\s*[-]*\d+\.\d+\s+MEA/) {
	    $ss = $1;
	}
    }
    close(FILE);

    #last one
    print STO "# STOCKHOLM 1.0\n\n";
    if ($extra =~ /\S+/) {
	print STO "#=GS $name$extra\n\n";
    }
    print STO "$name         $seq\n";
    print STO "#=GR $name SS $ss\n";
    print STO "\/\/\n";

    close(STO);

    return $viennasto;
}
```
