## Supplementary figures and images for "What does it take to learn the rules of RNA base pairing? A lot less than you may think"

### Figure_R1.pdf

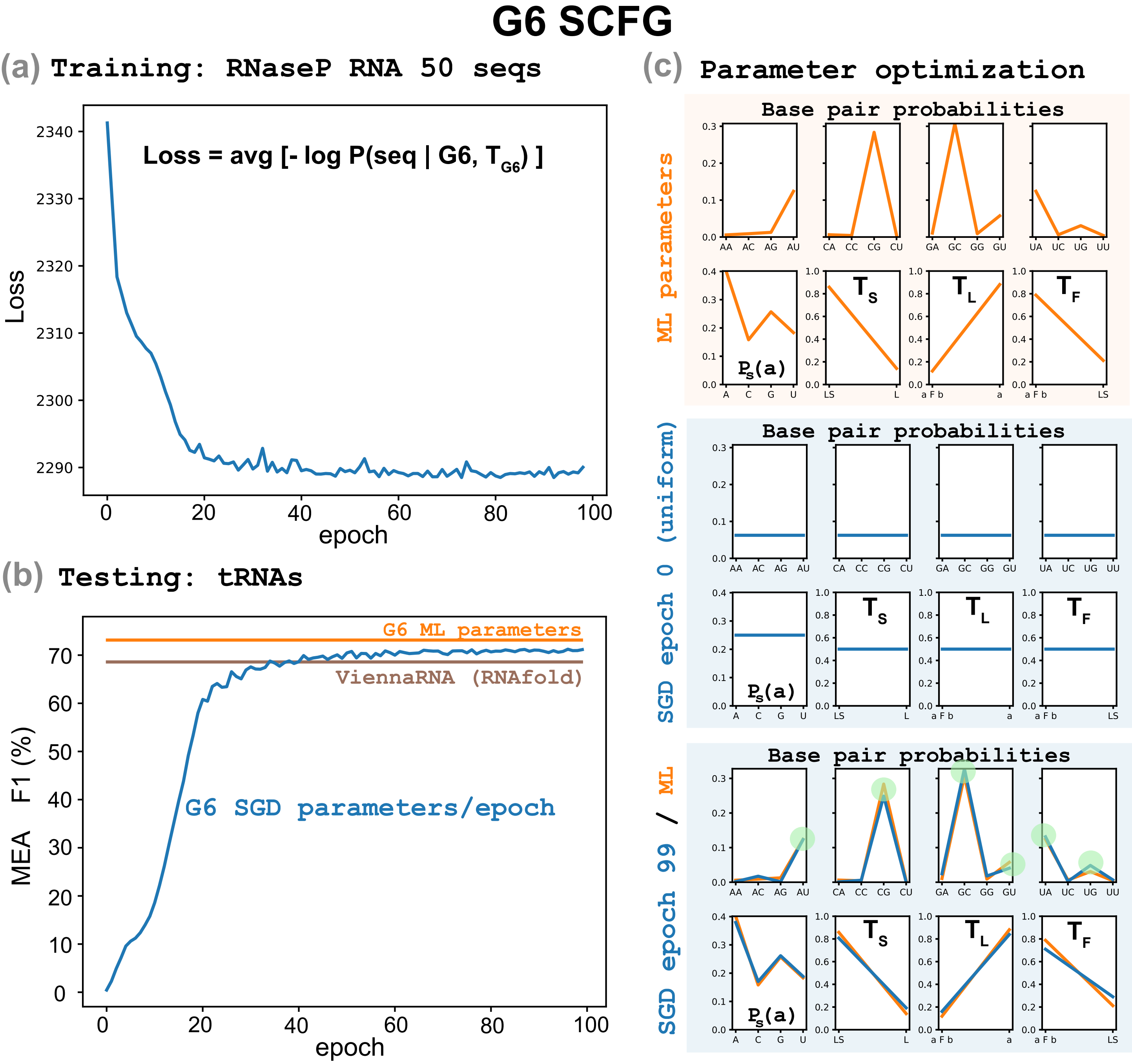

### Figure_R2.pdf

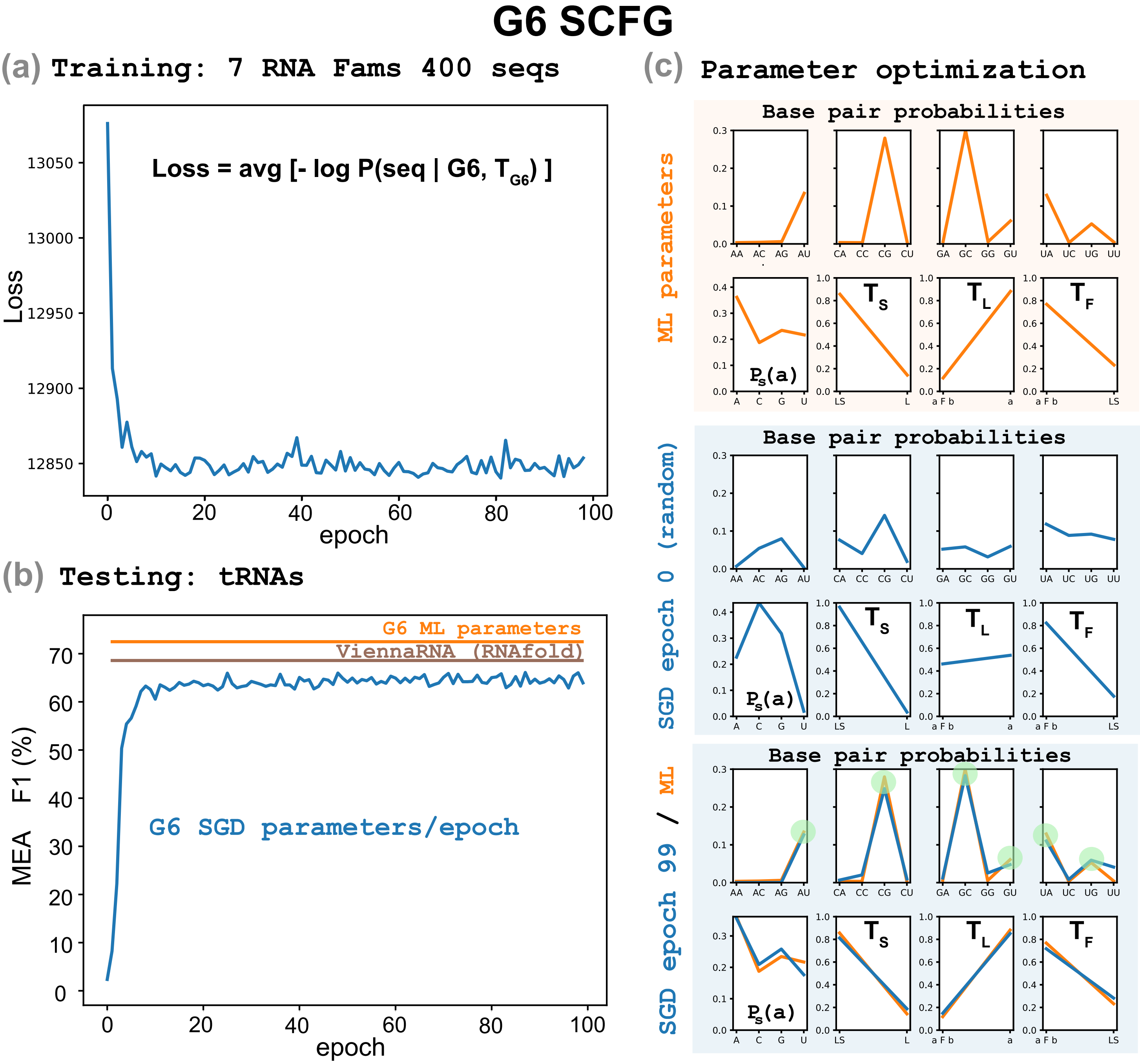

### Figure_R3.pdf

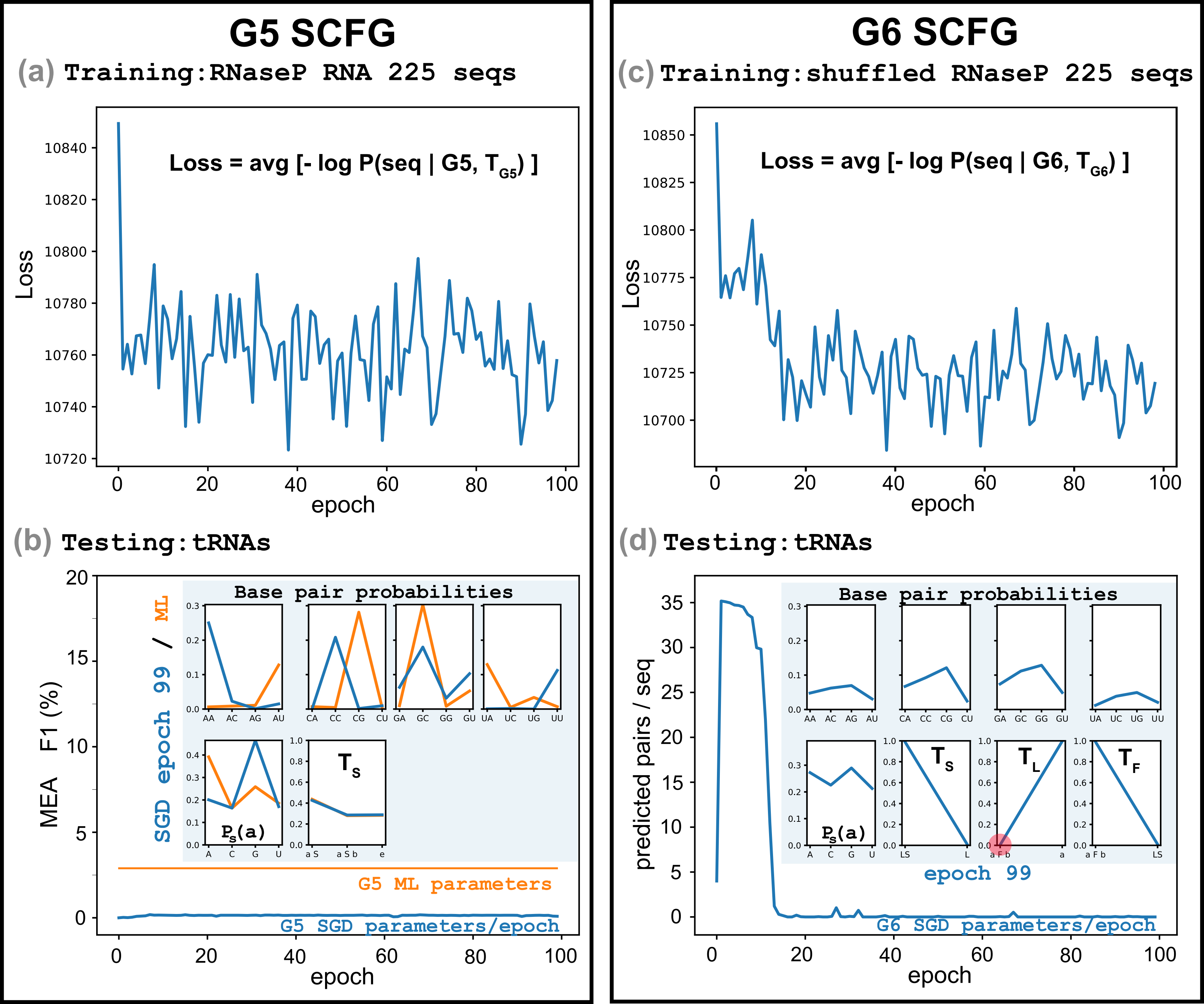

### Figure_R4.pdf

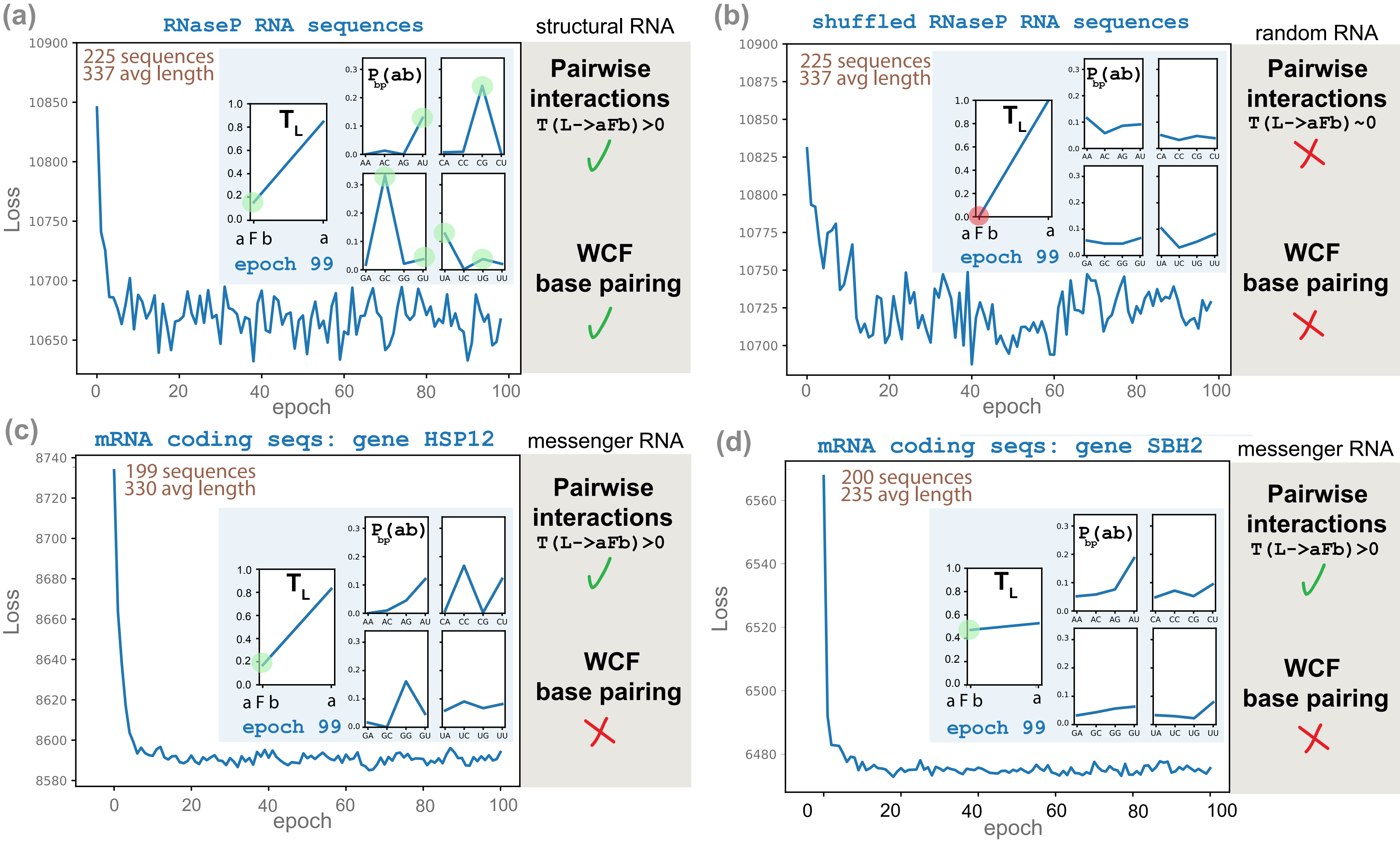

### Figure_S1.pdf

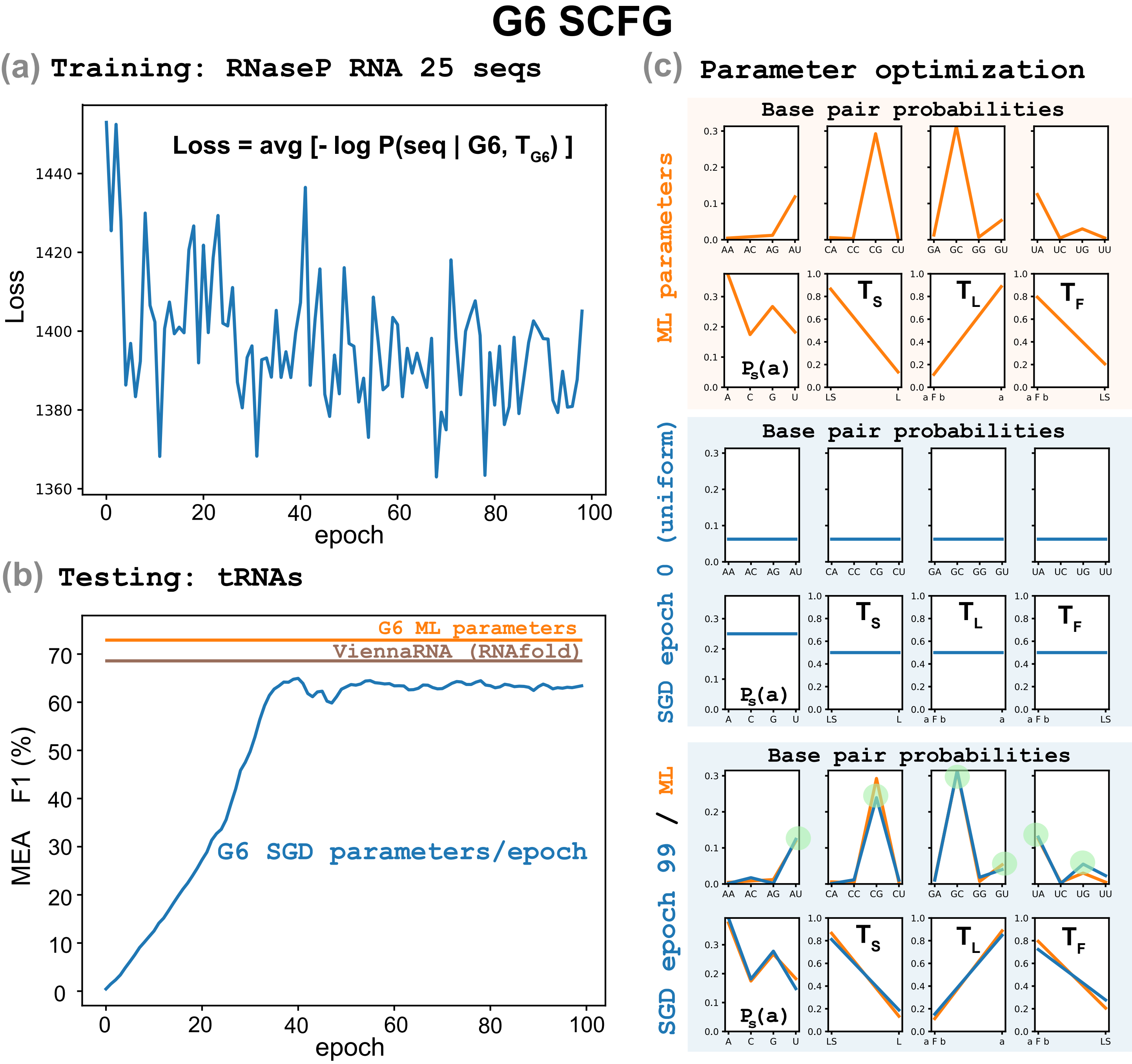

### g6_params_i48.pdf

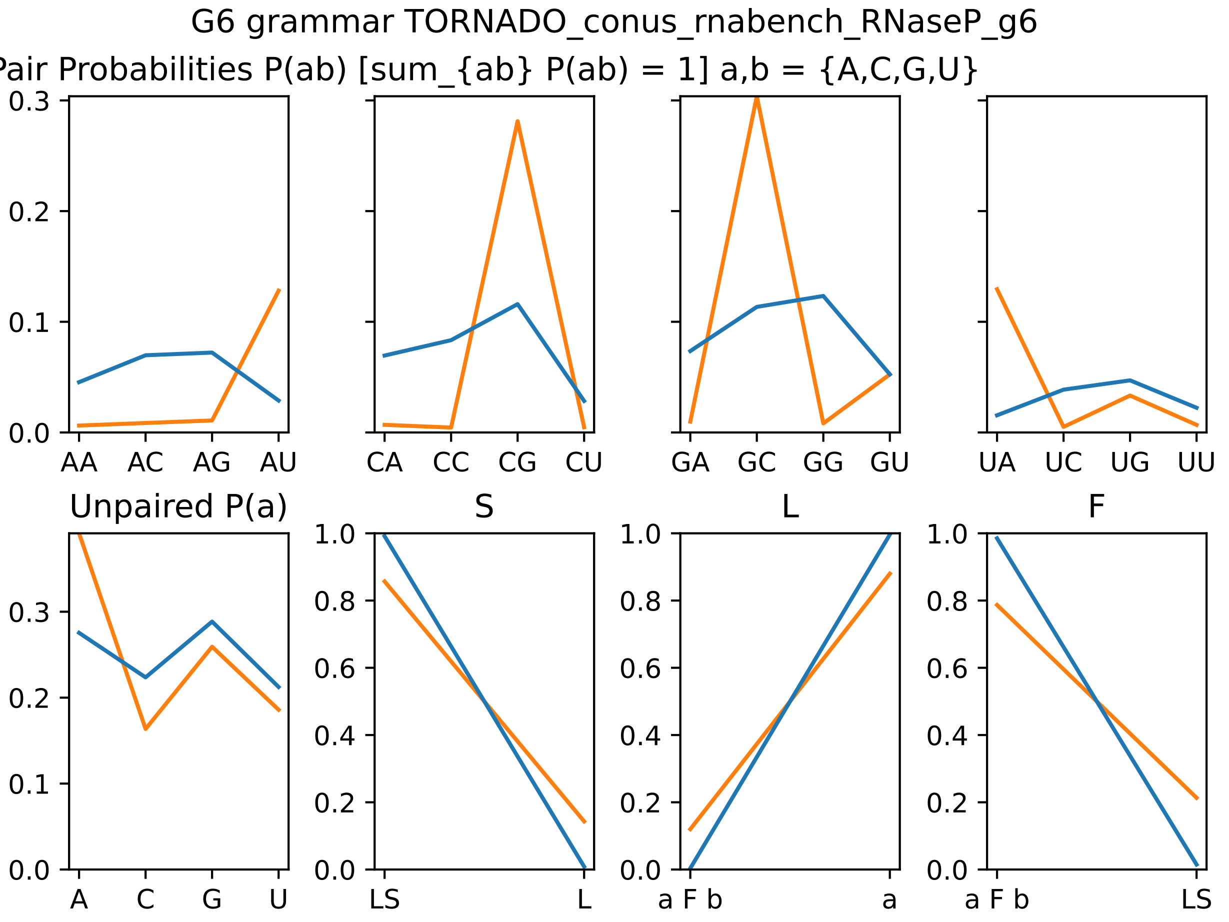

### g6_params_i49.pdf

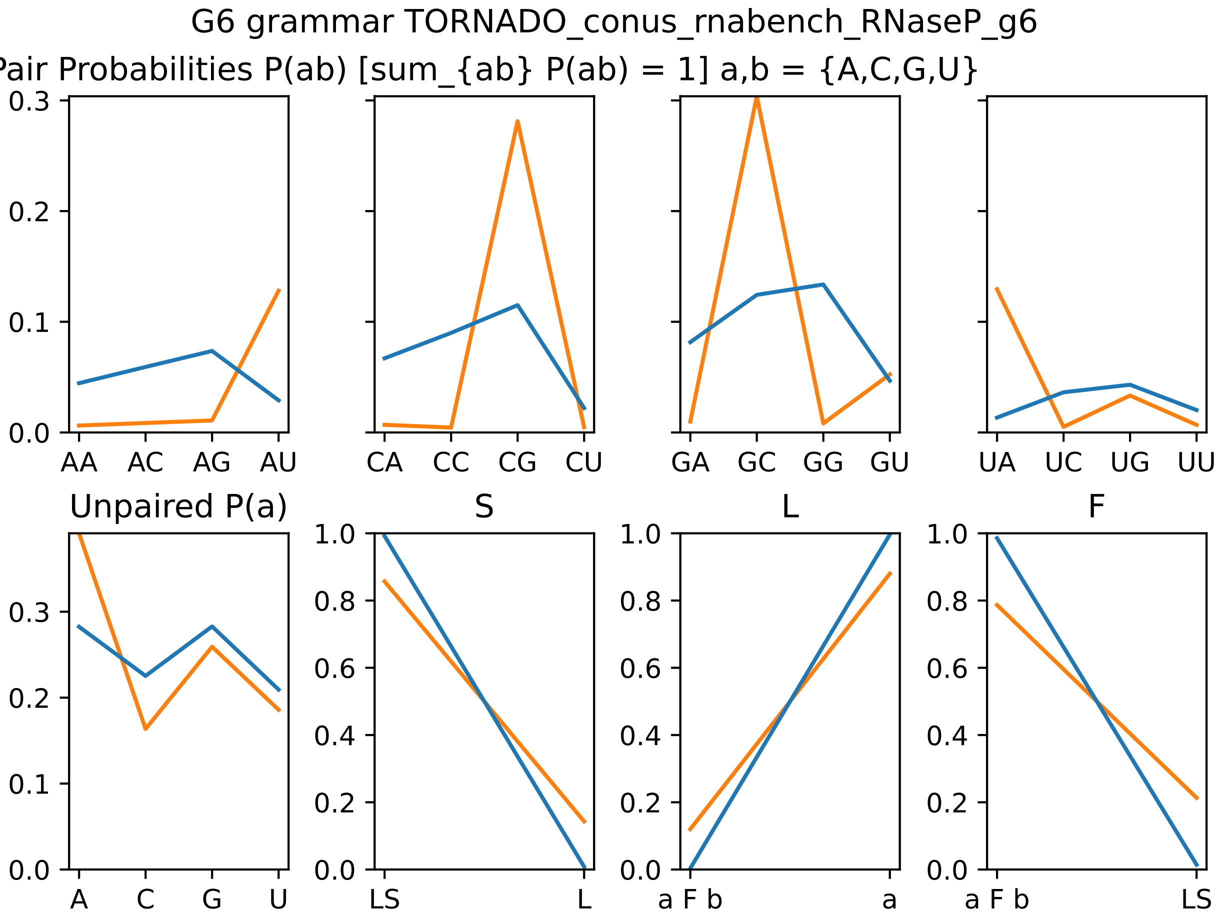

### g6_params_i60.pdf

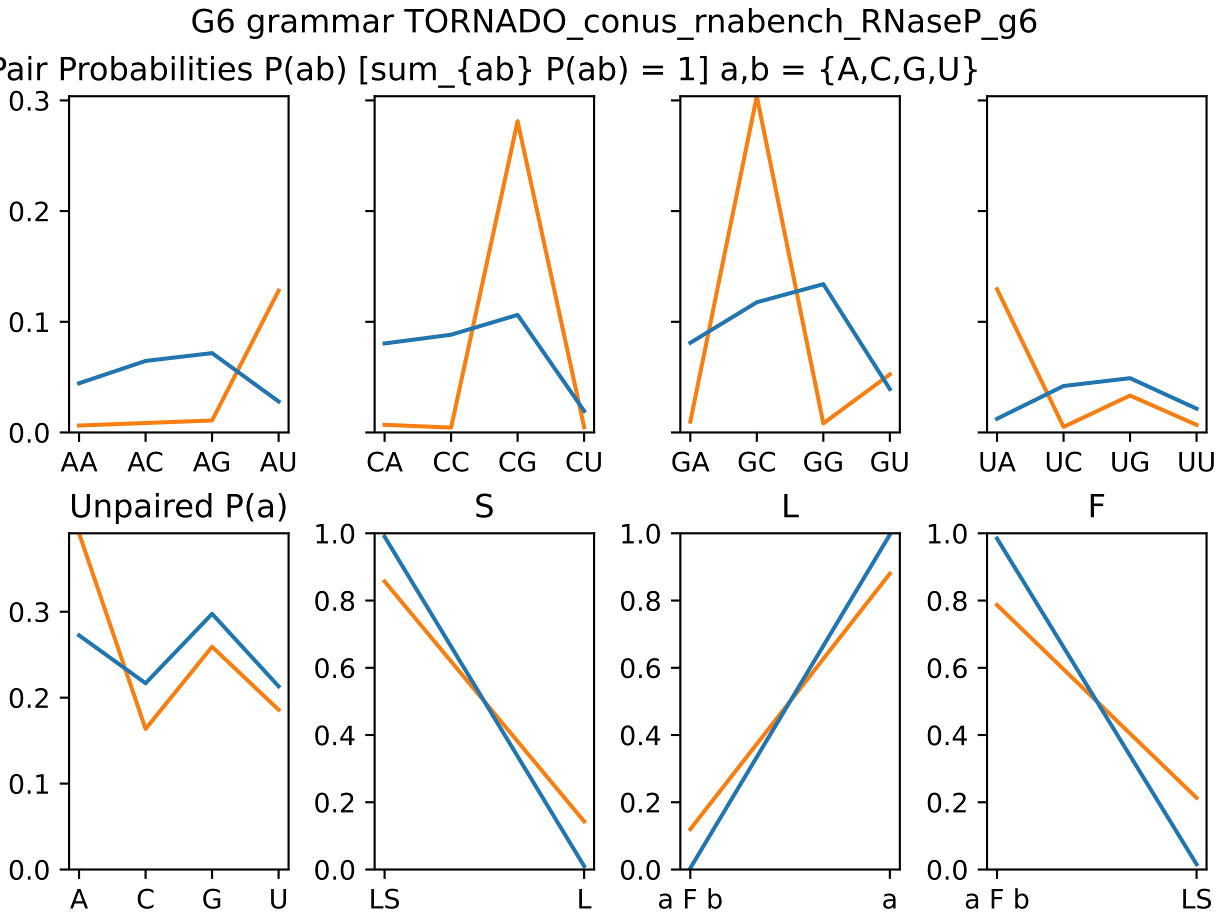

### g6_params_i61.pdf

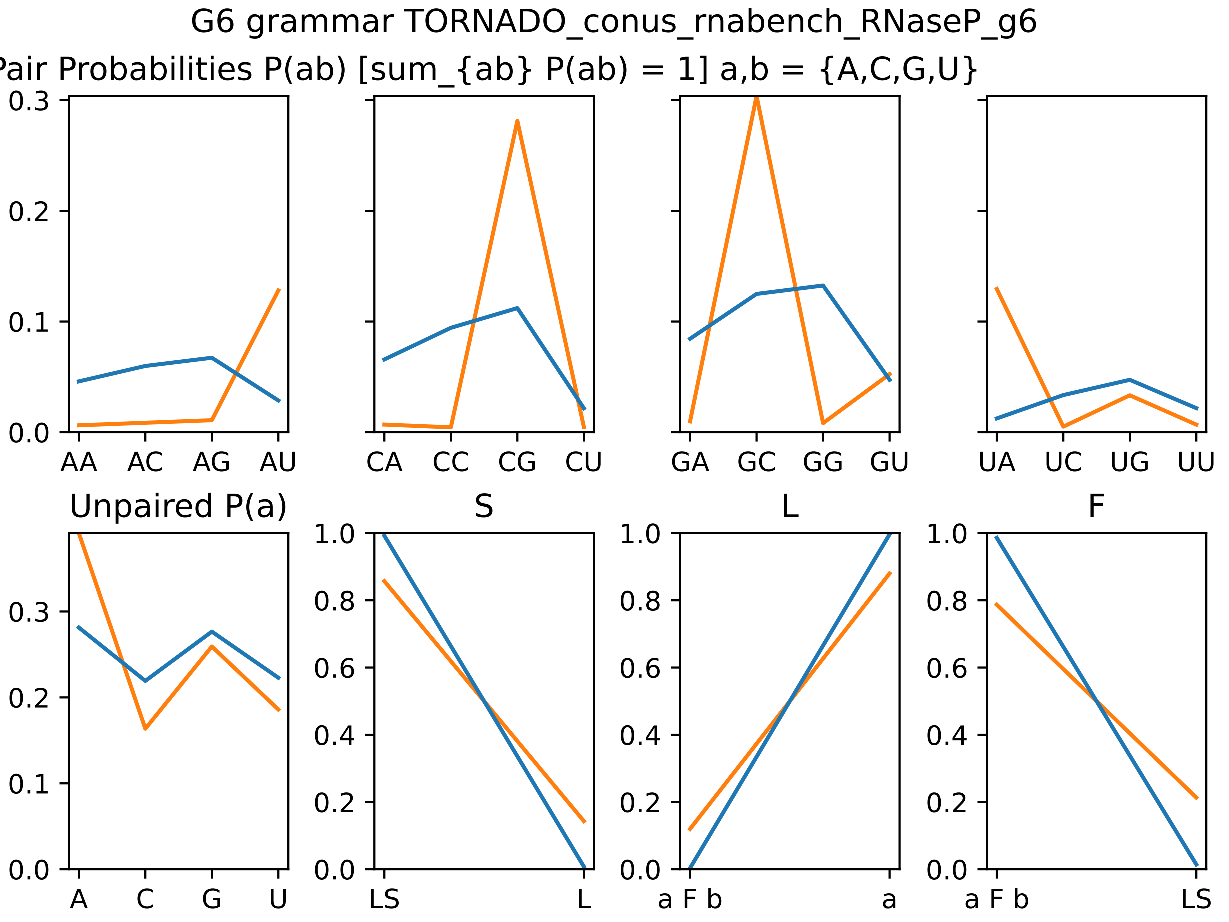

### g6_params_i62.pdf

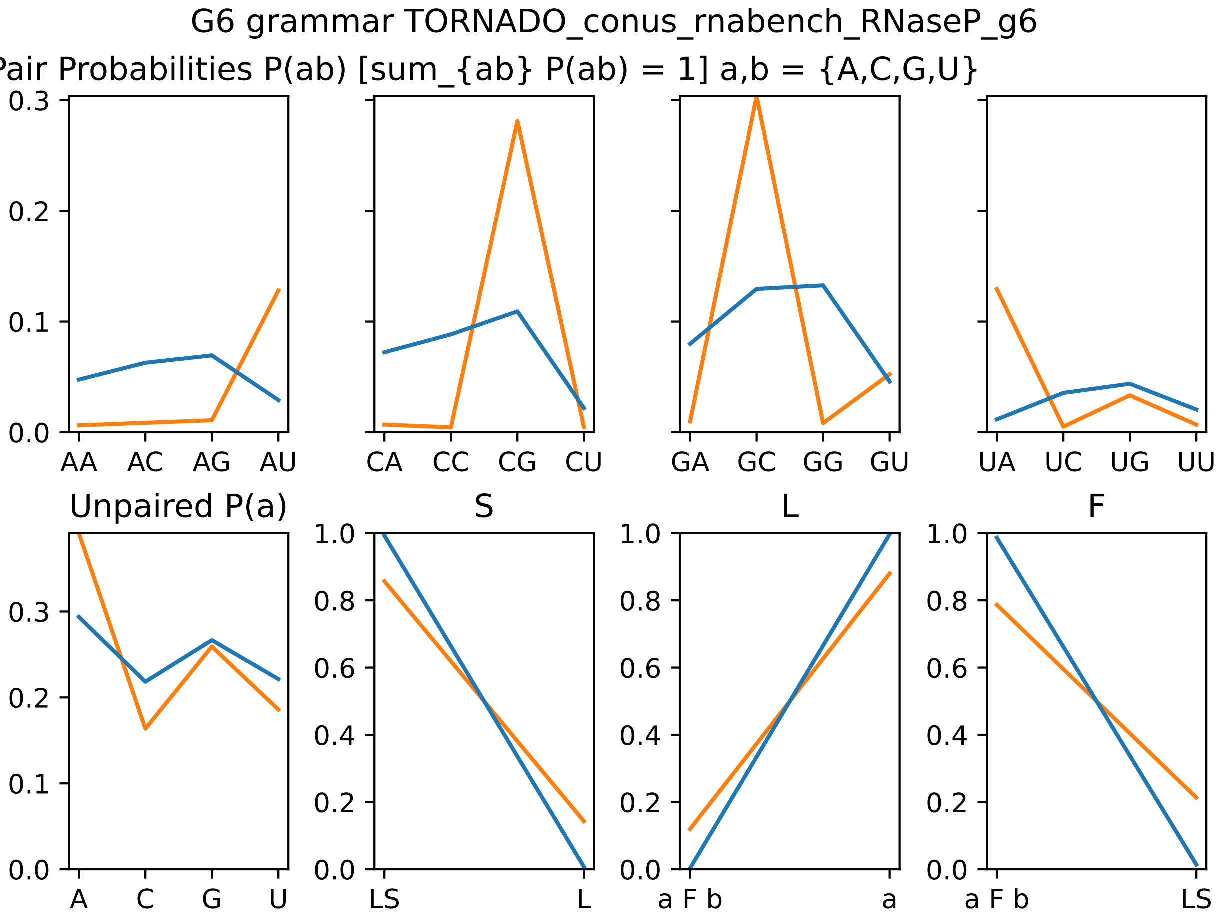

### g6_params_i63.pdf

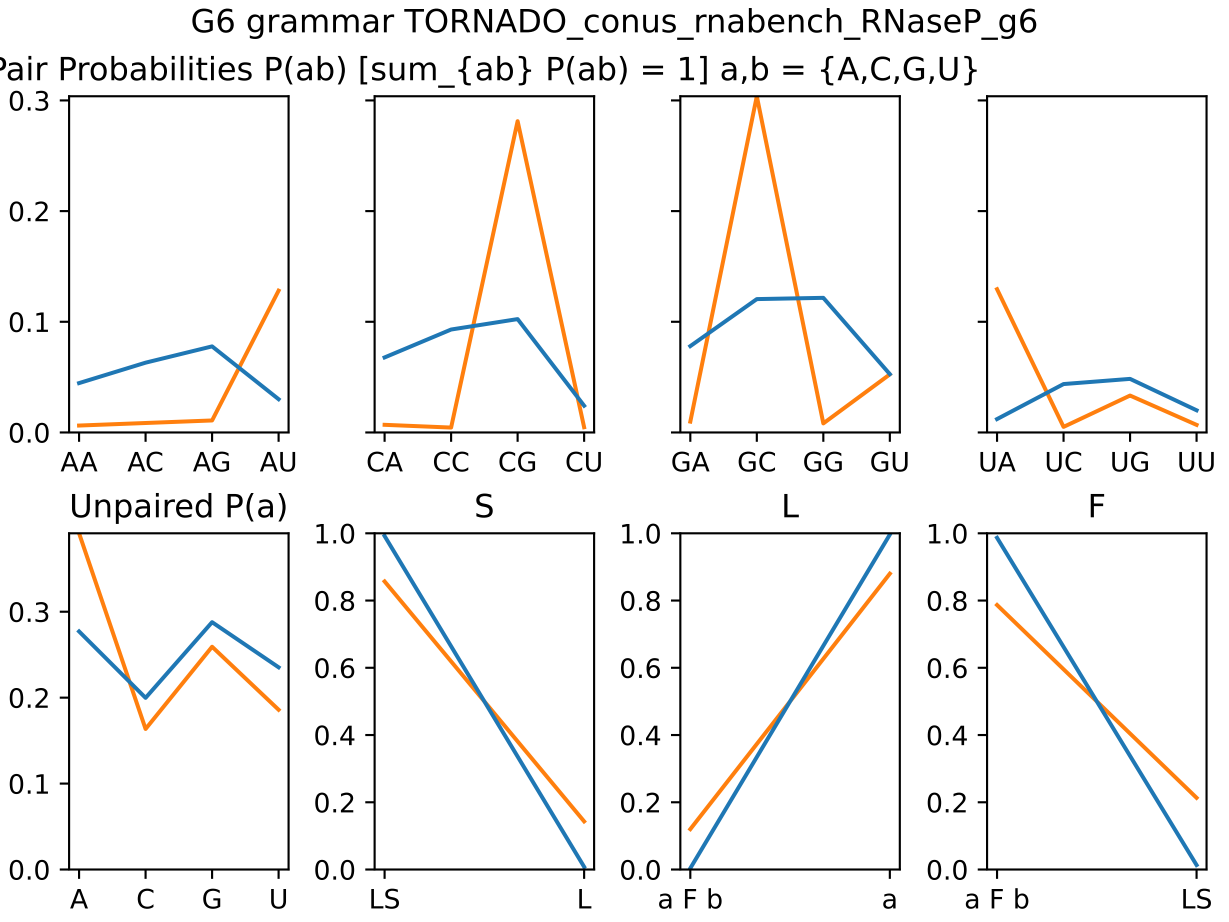

### g6_params_i64.pdf

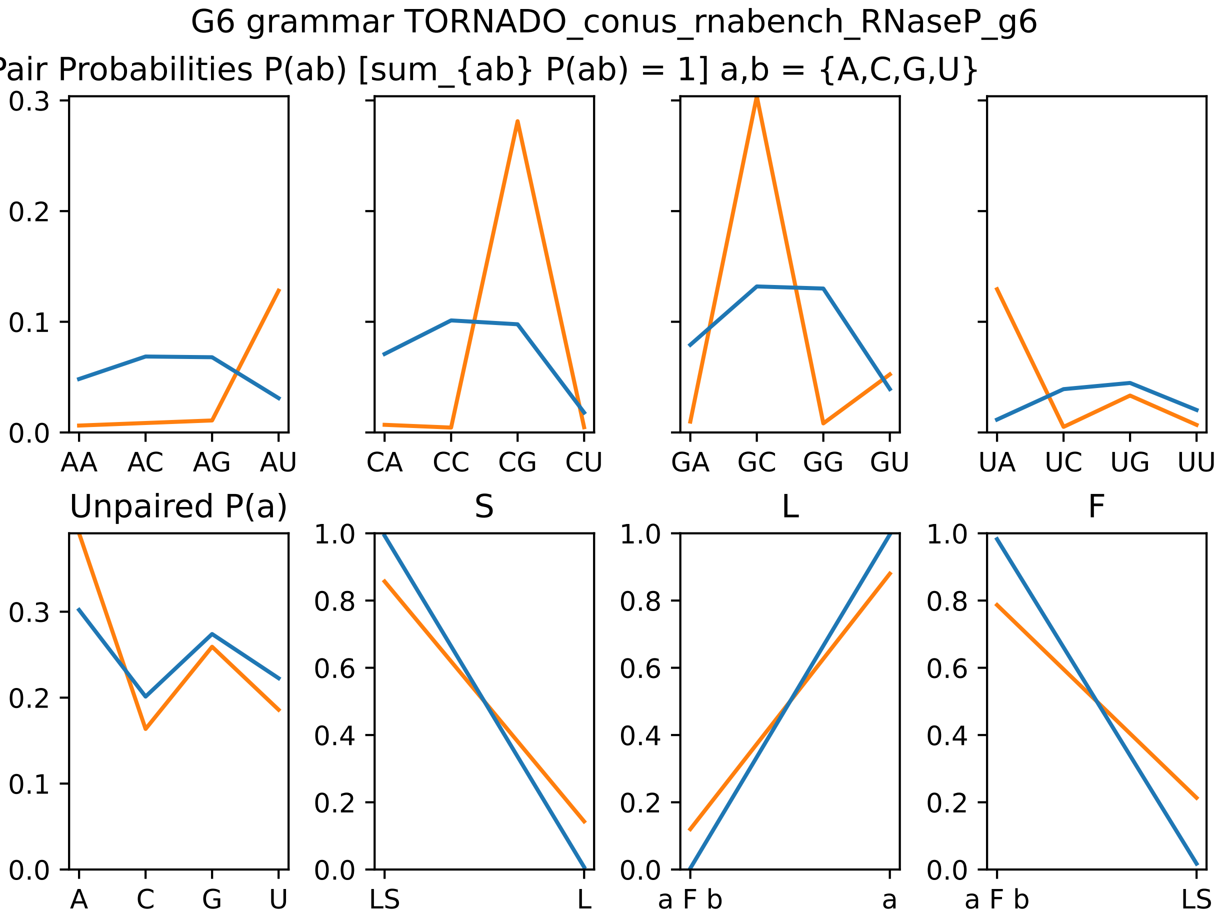

### g6_params_i70.pdf

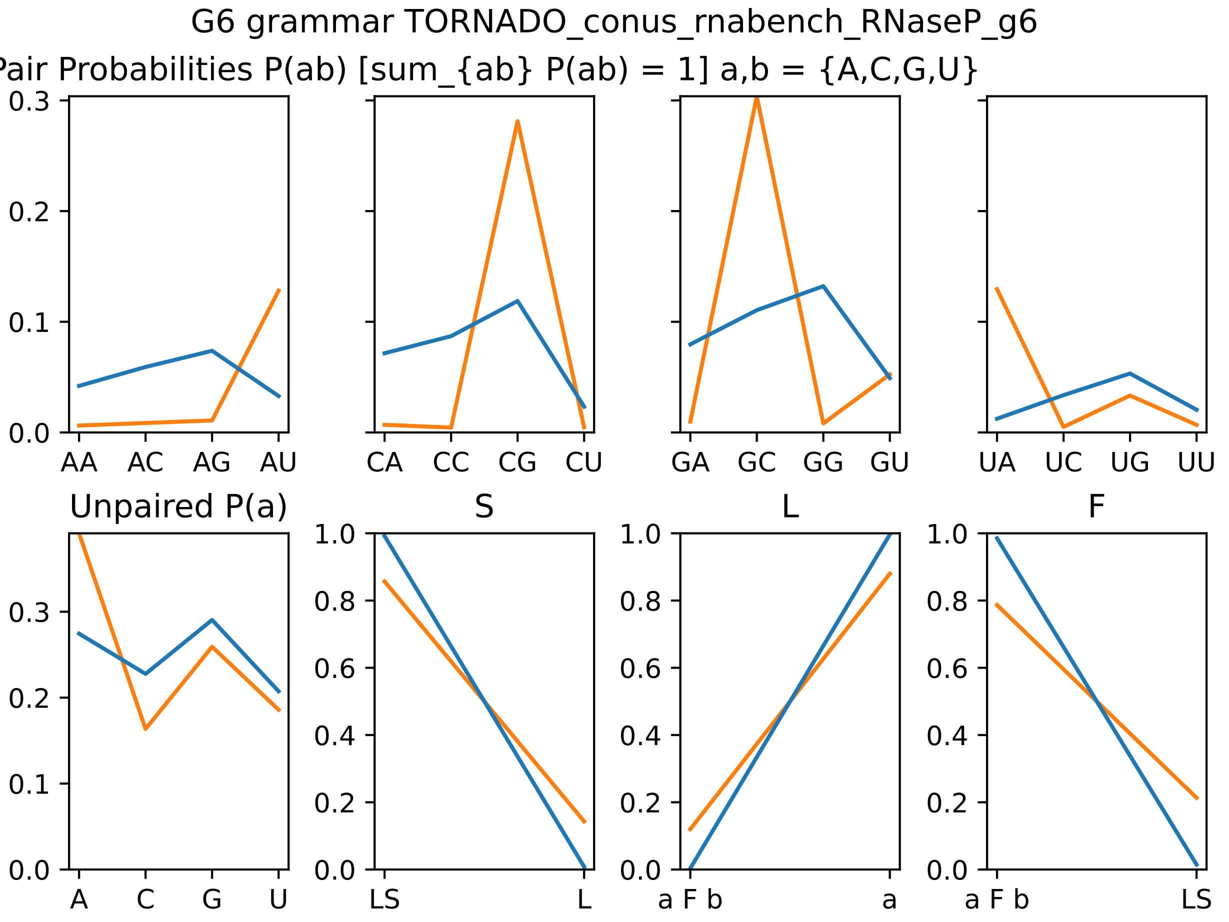

### g6_params_i74.pdf

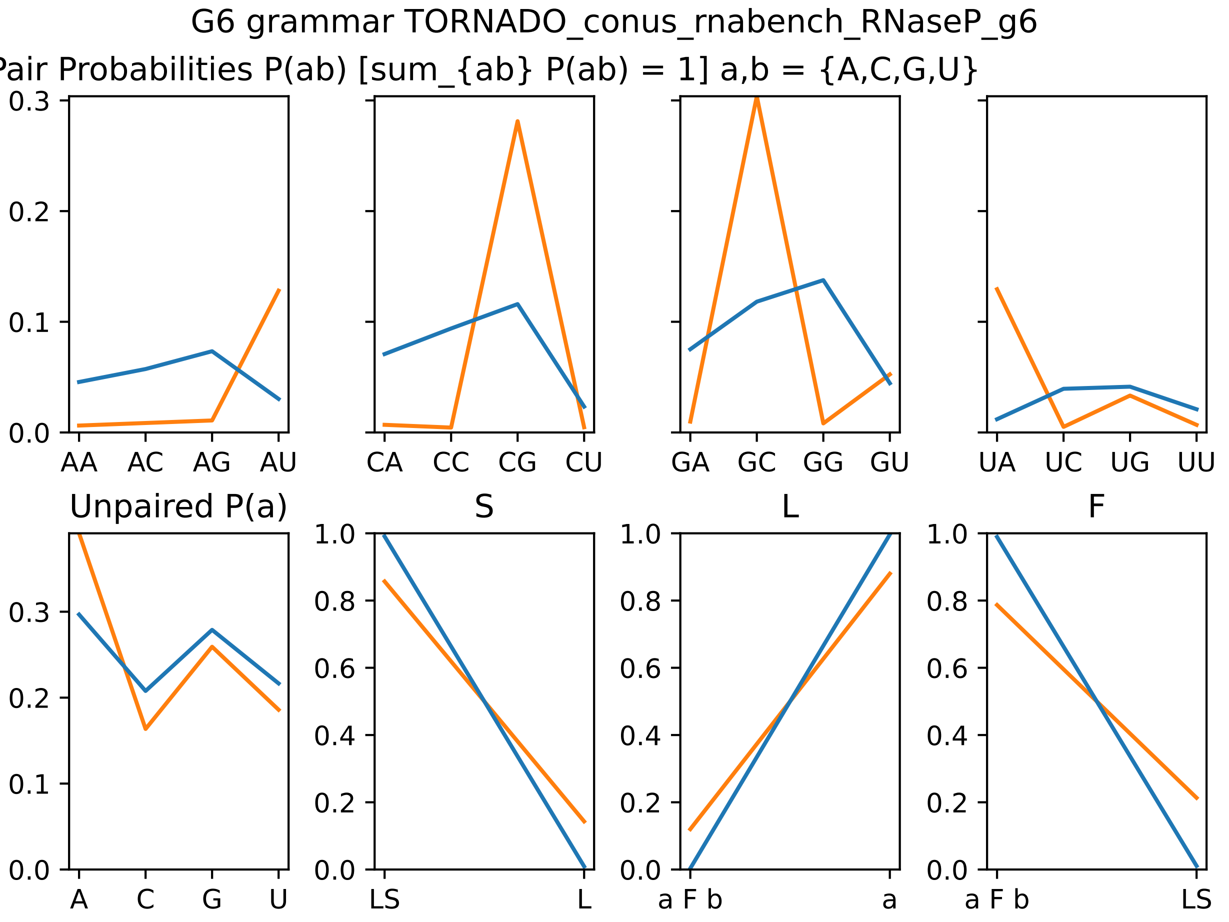

### g6_params_i75.pdf

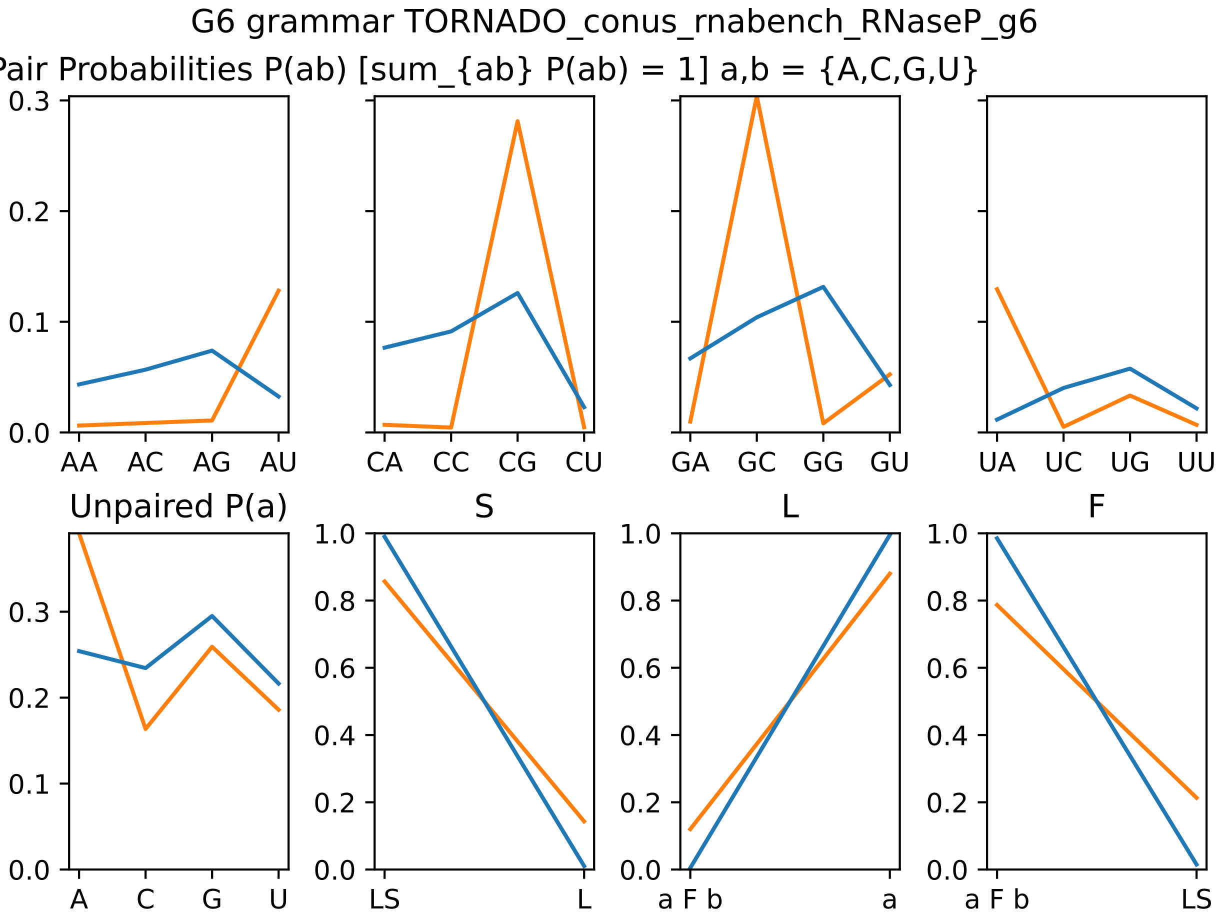

### g6_params_i76.pdf

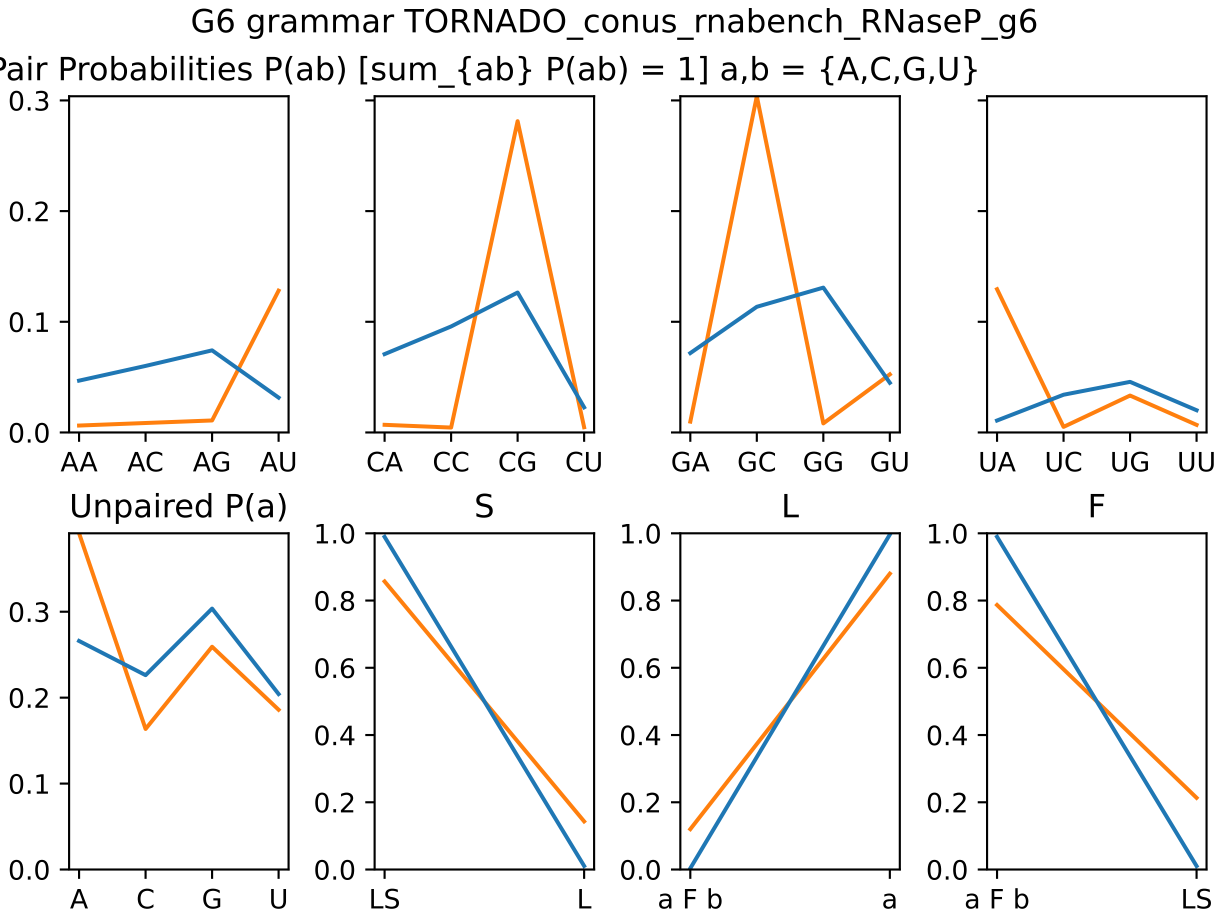

### g6_params_i77.pdf

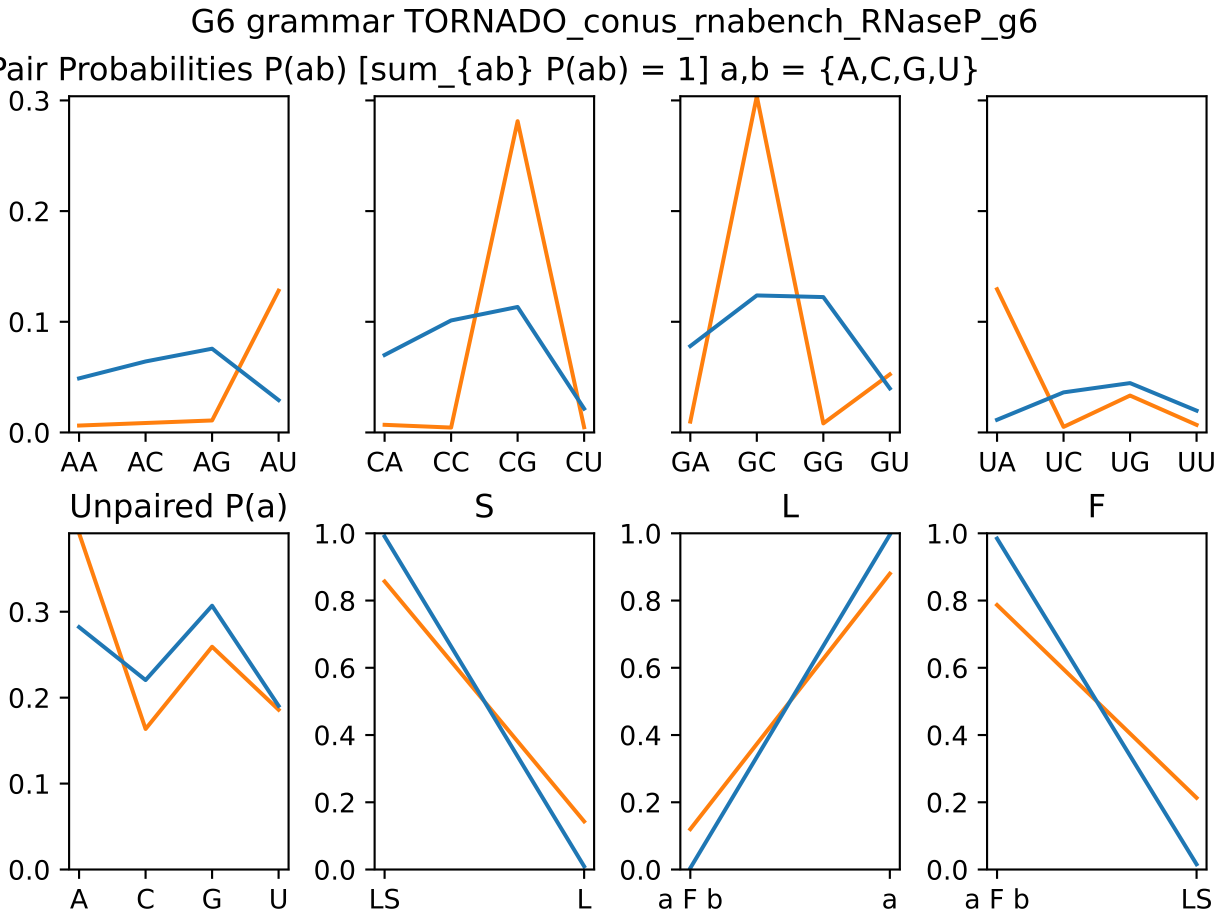

### g6_params_i88.pdf

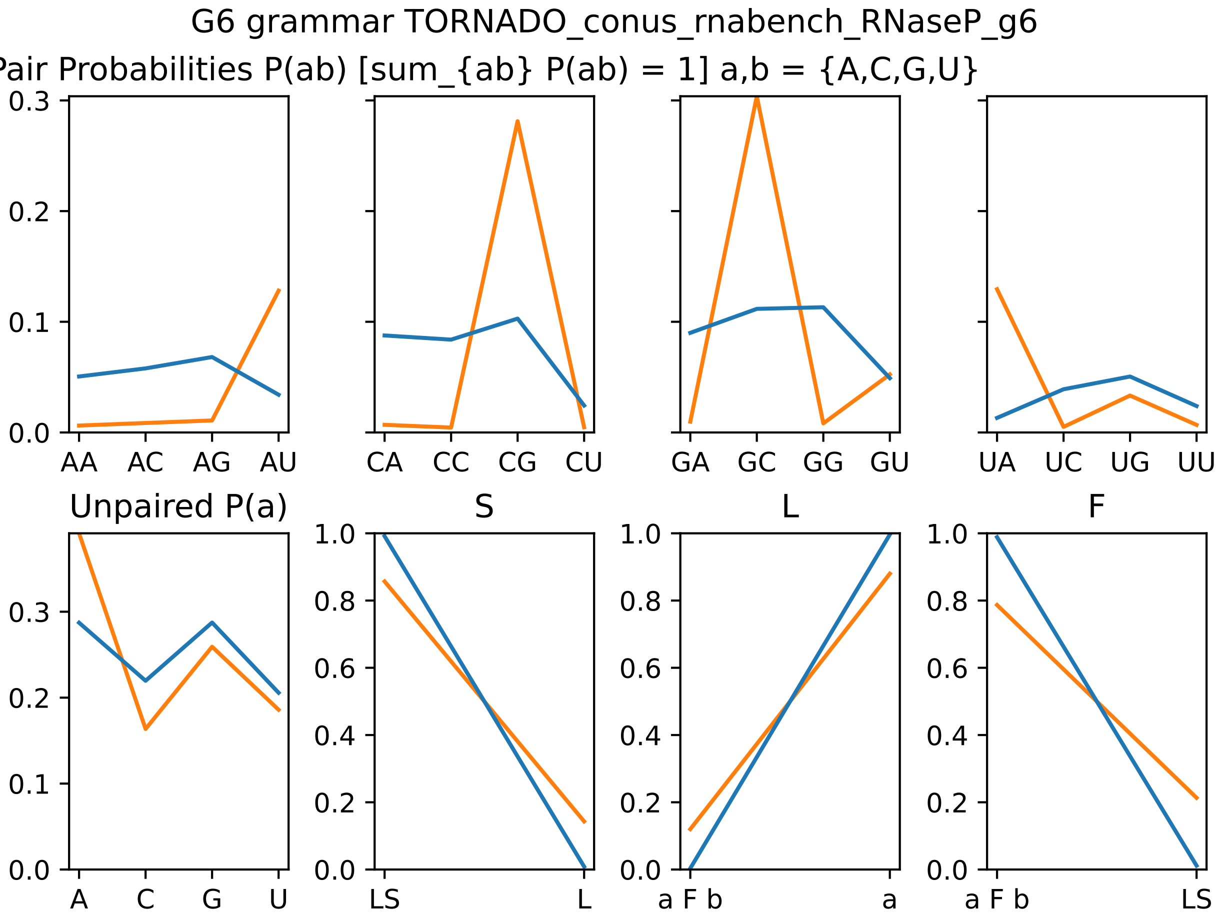

### g6_params_i89.pdf

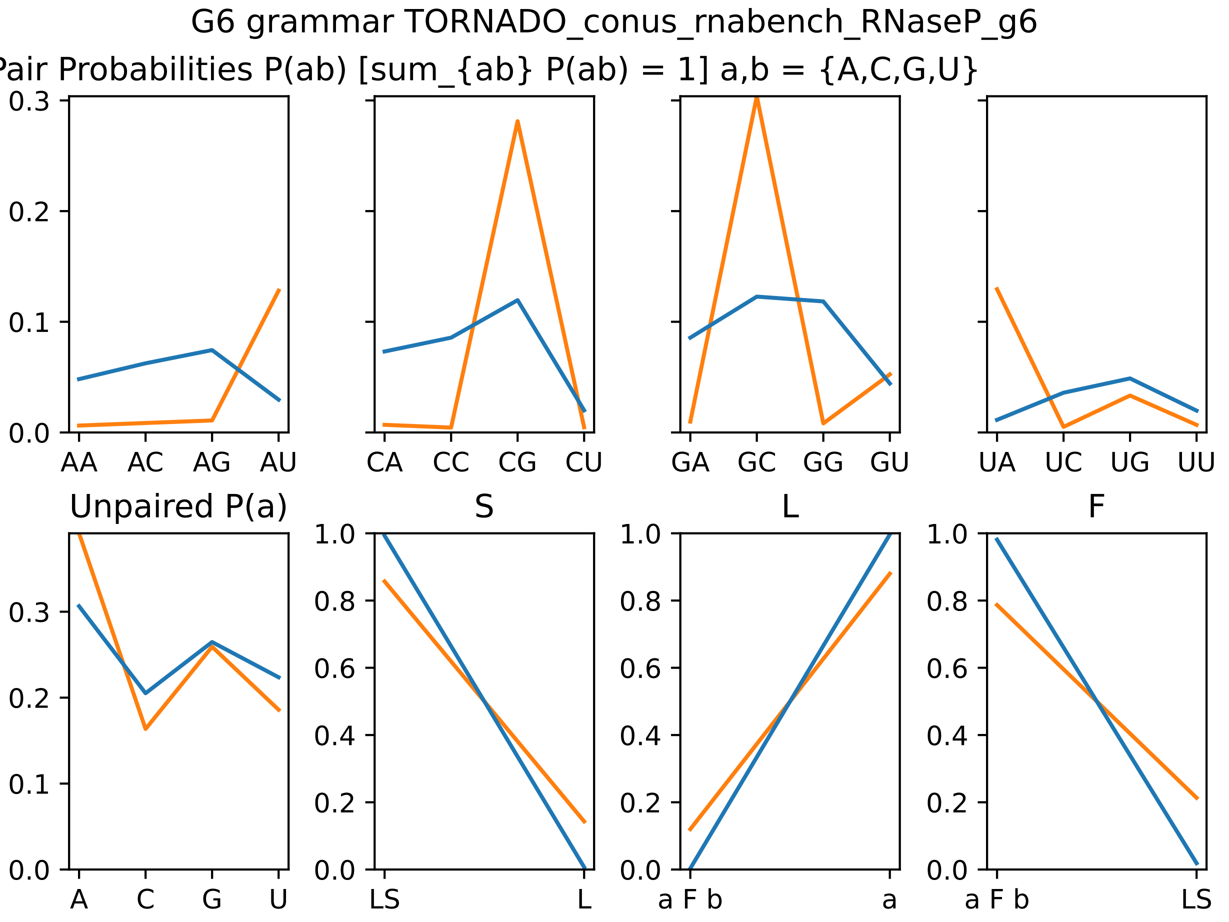

### losses_i40.pdf

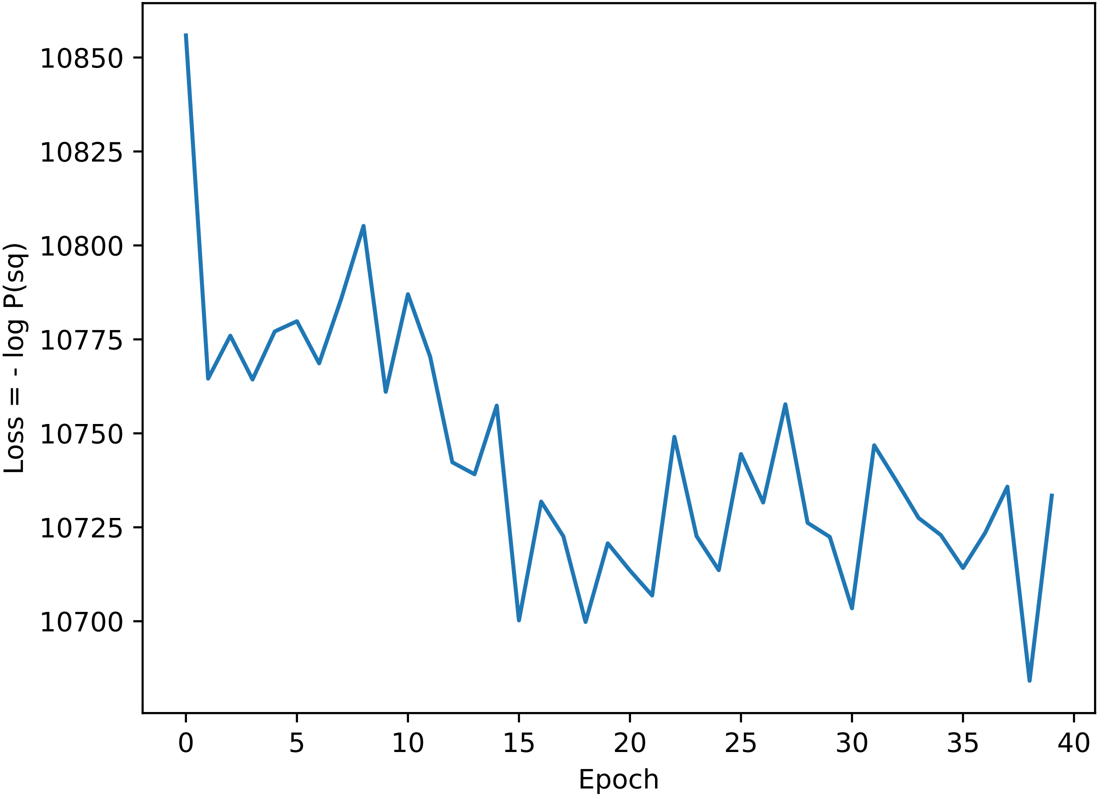

### losses_i41.pdf

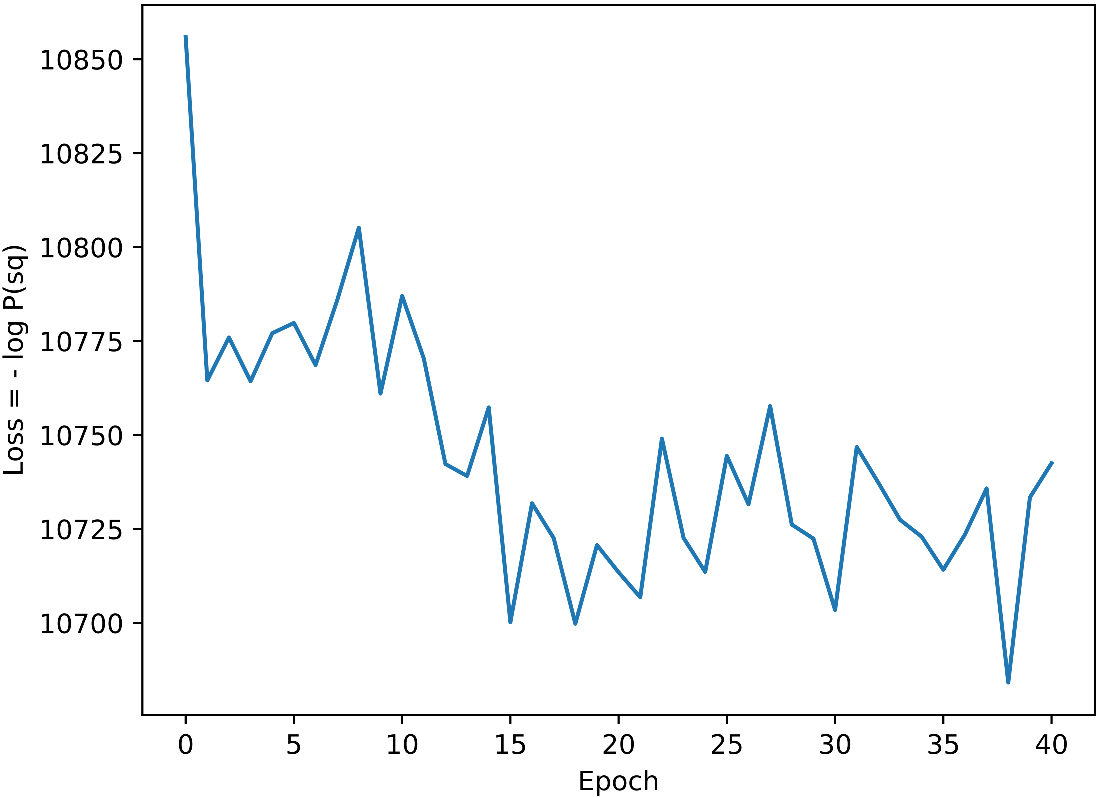

### losses_i42.pdf

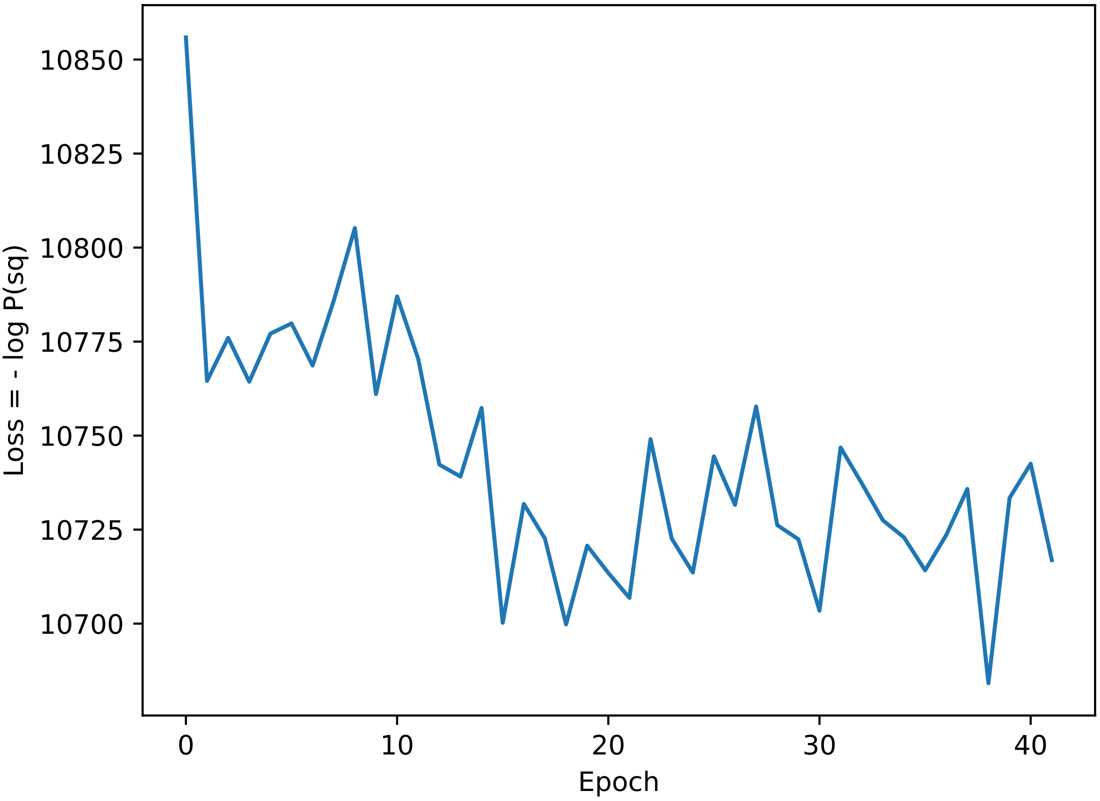

### losses_i43.pdf

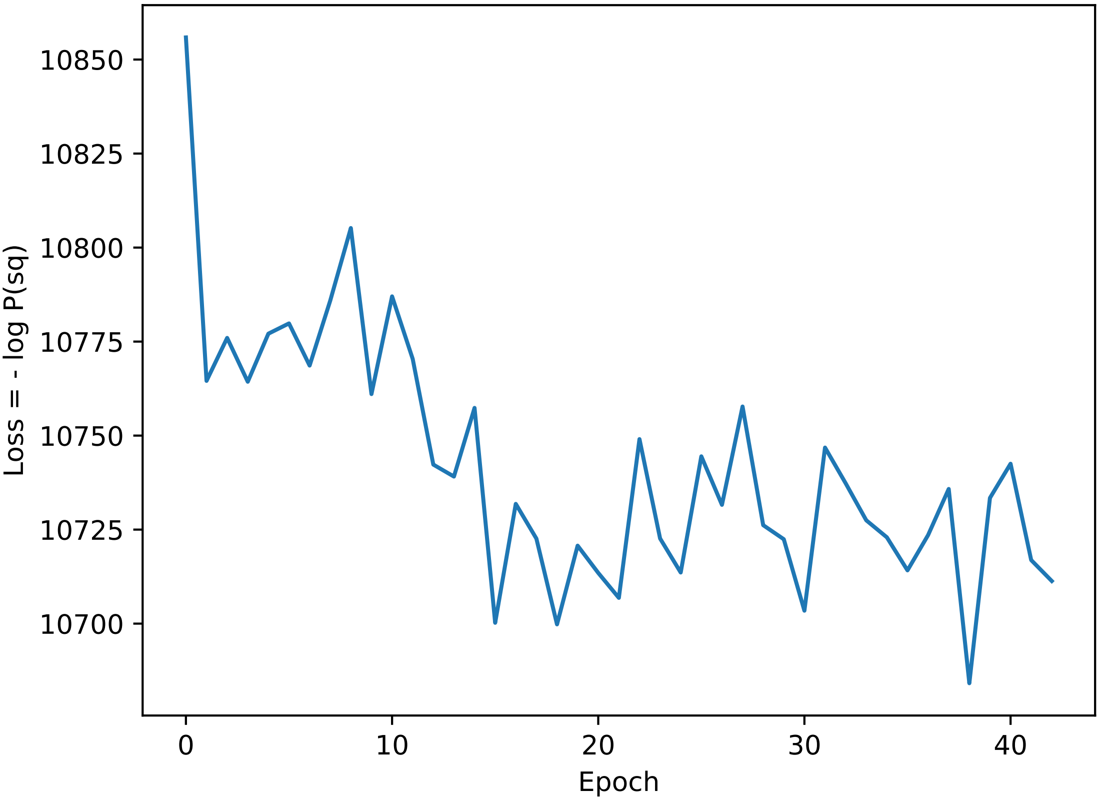

### losses_i54.pdf

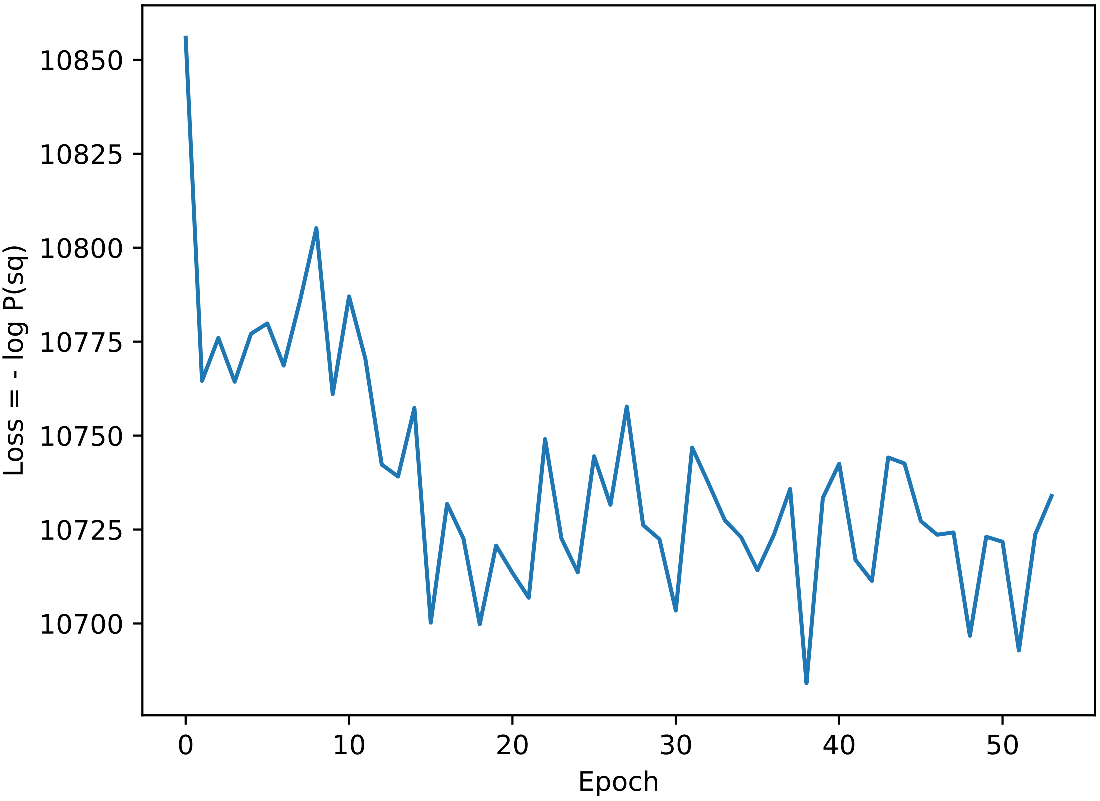

### losses_i55.pdf

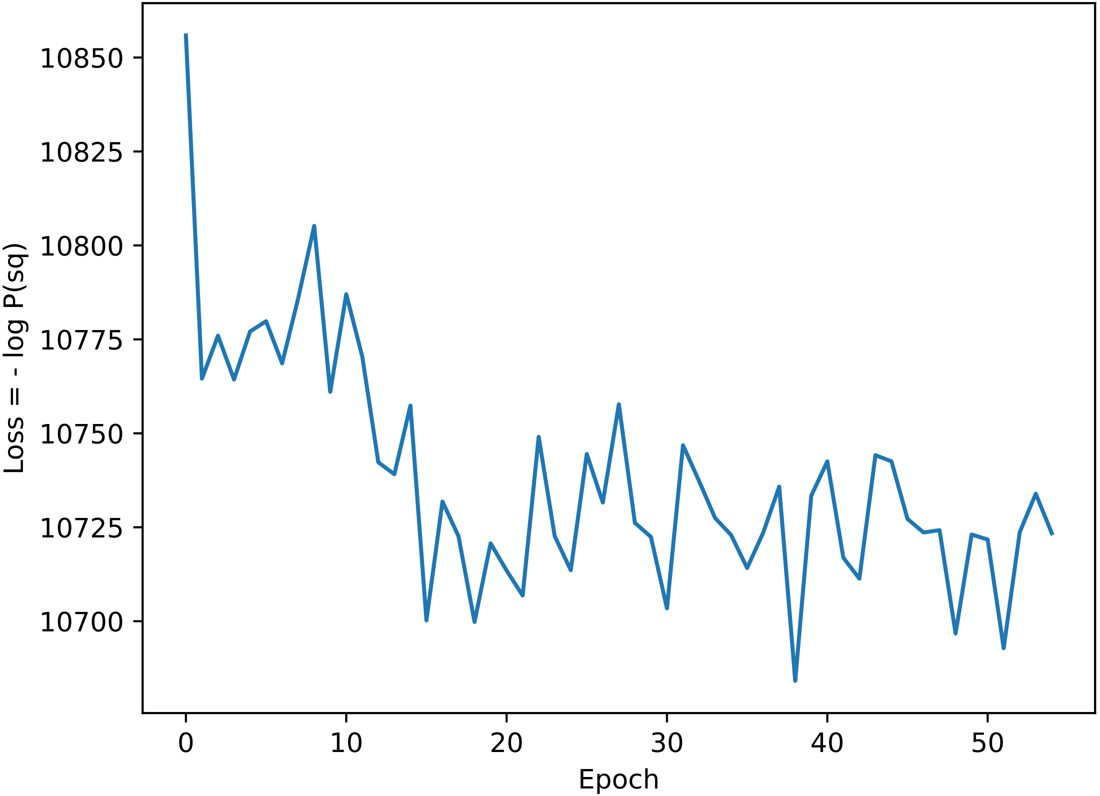

### losses_i56.pdf

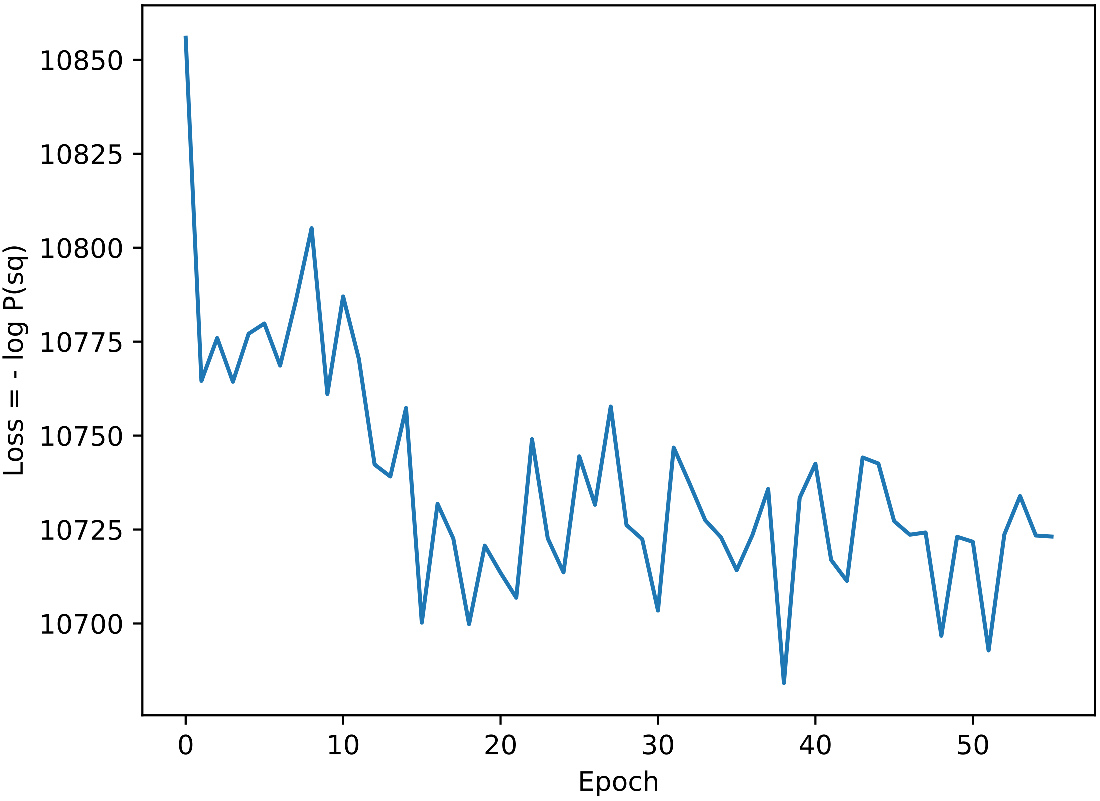

### losses_i57.pdf

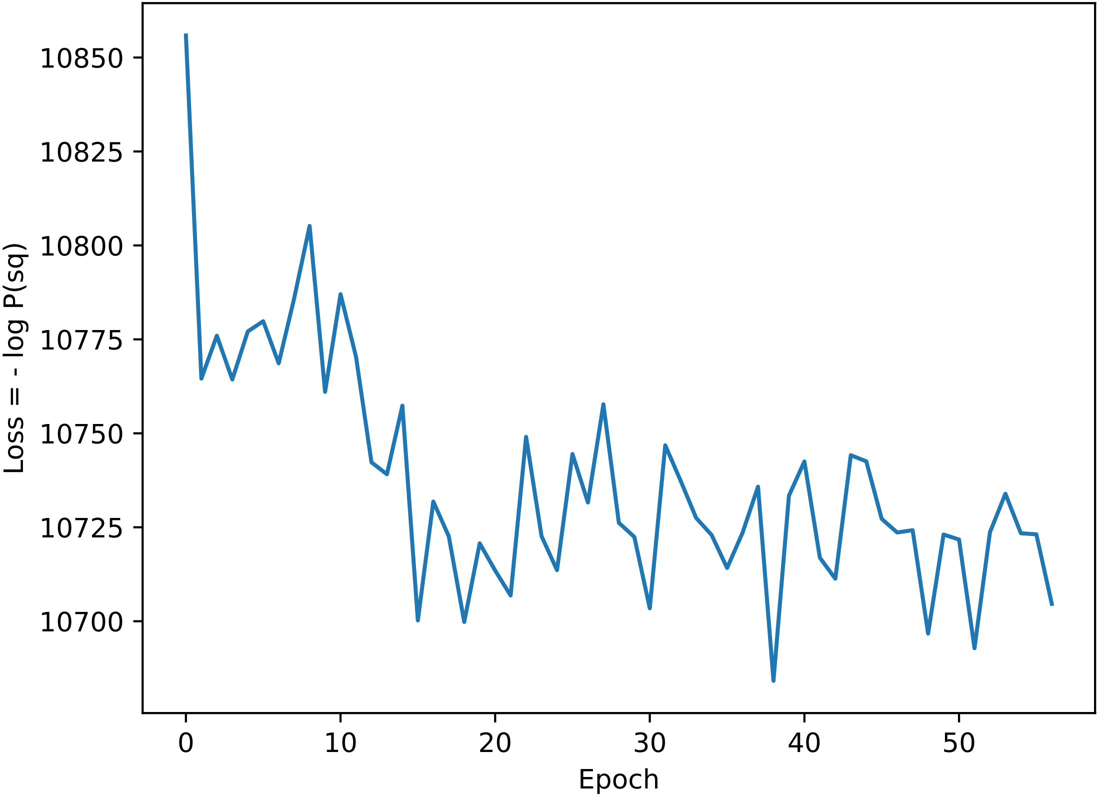

### losses_i68.pdf

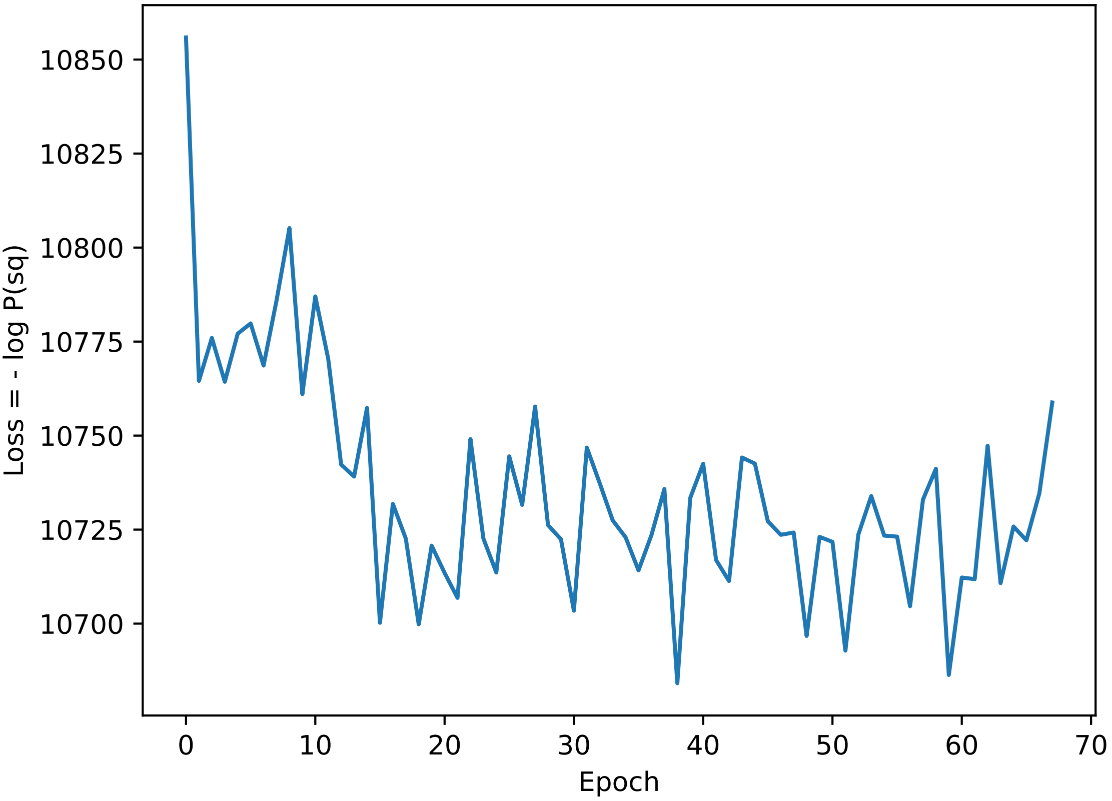

### losses_i69.pdf

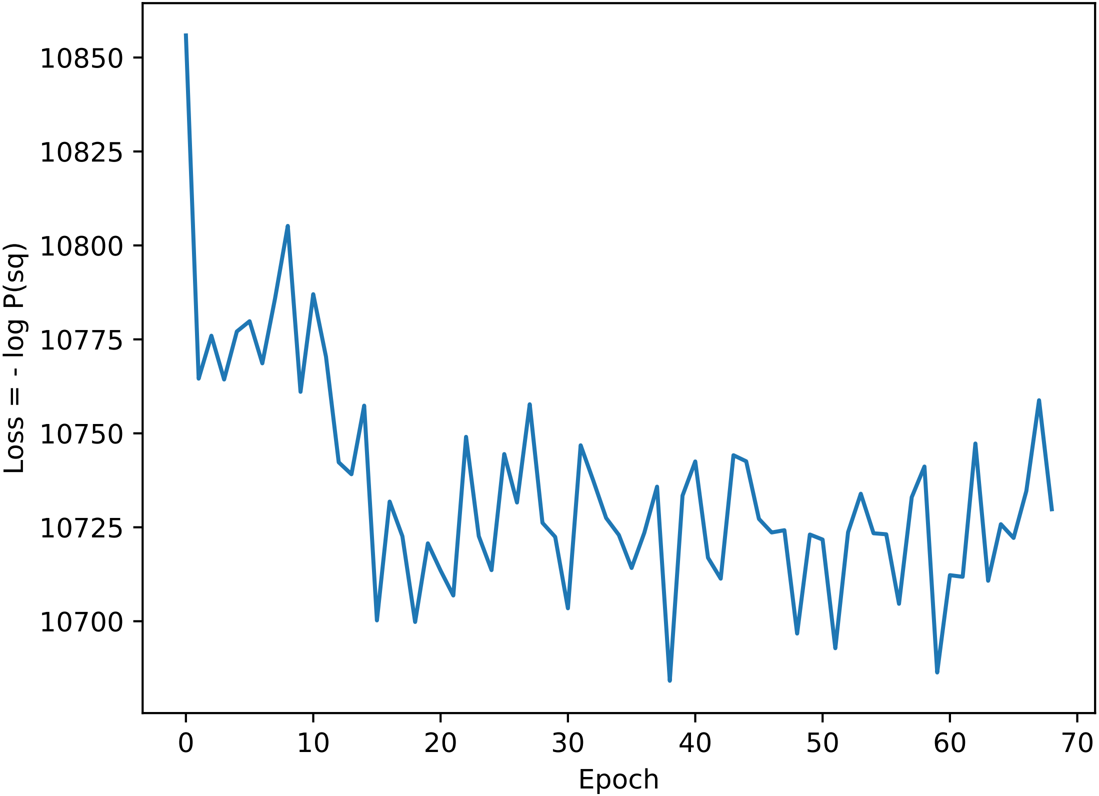

### losses_i80.pdf

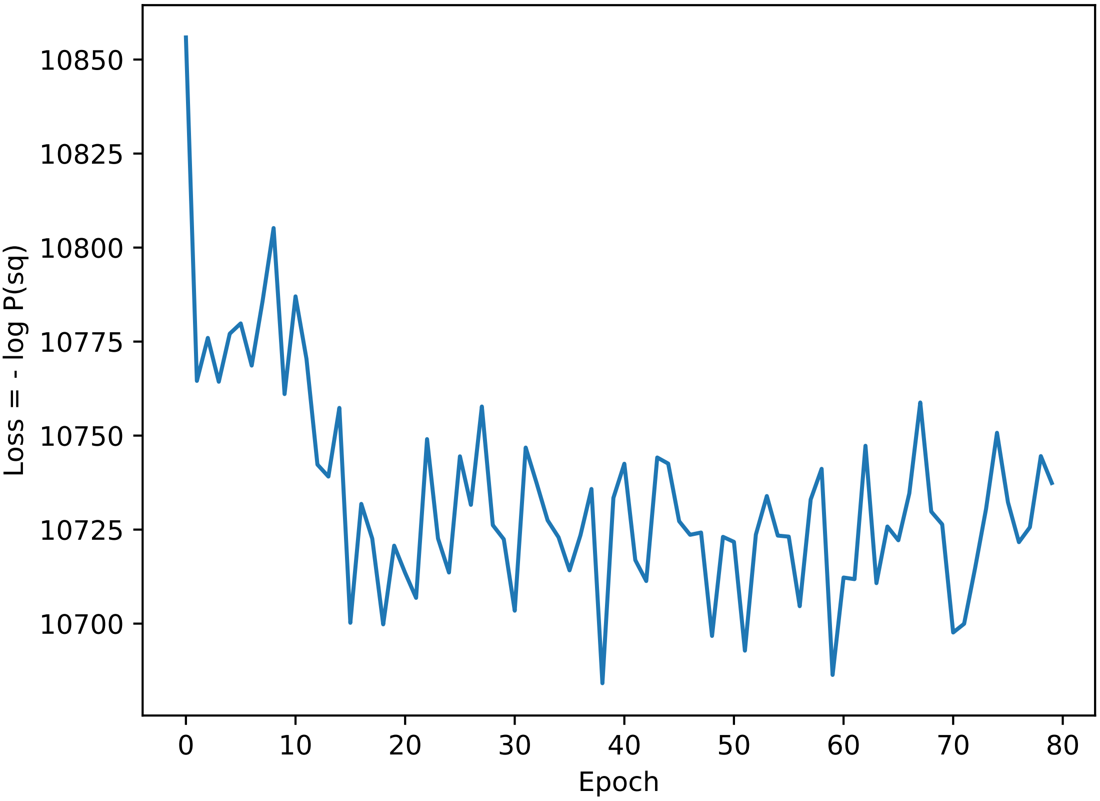

### losses_i81.pdf

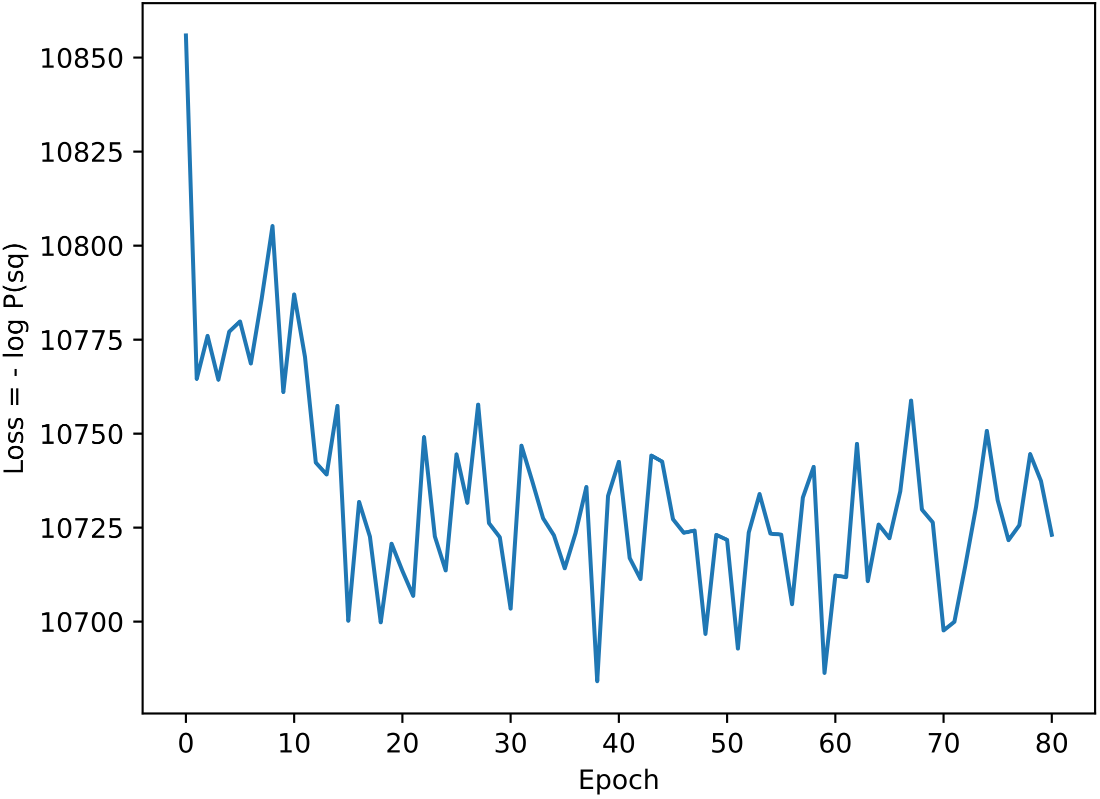

### losses_i82.pdf

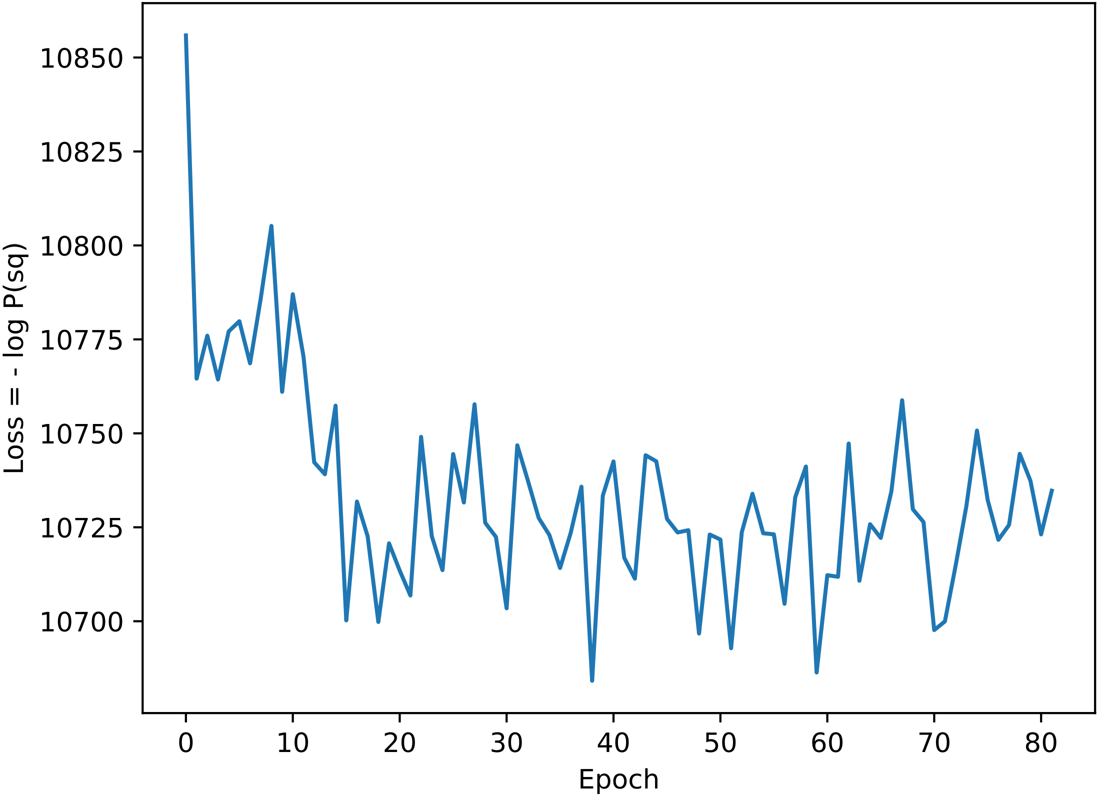

### losses_i83.pdf

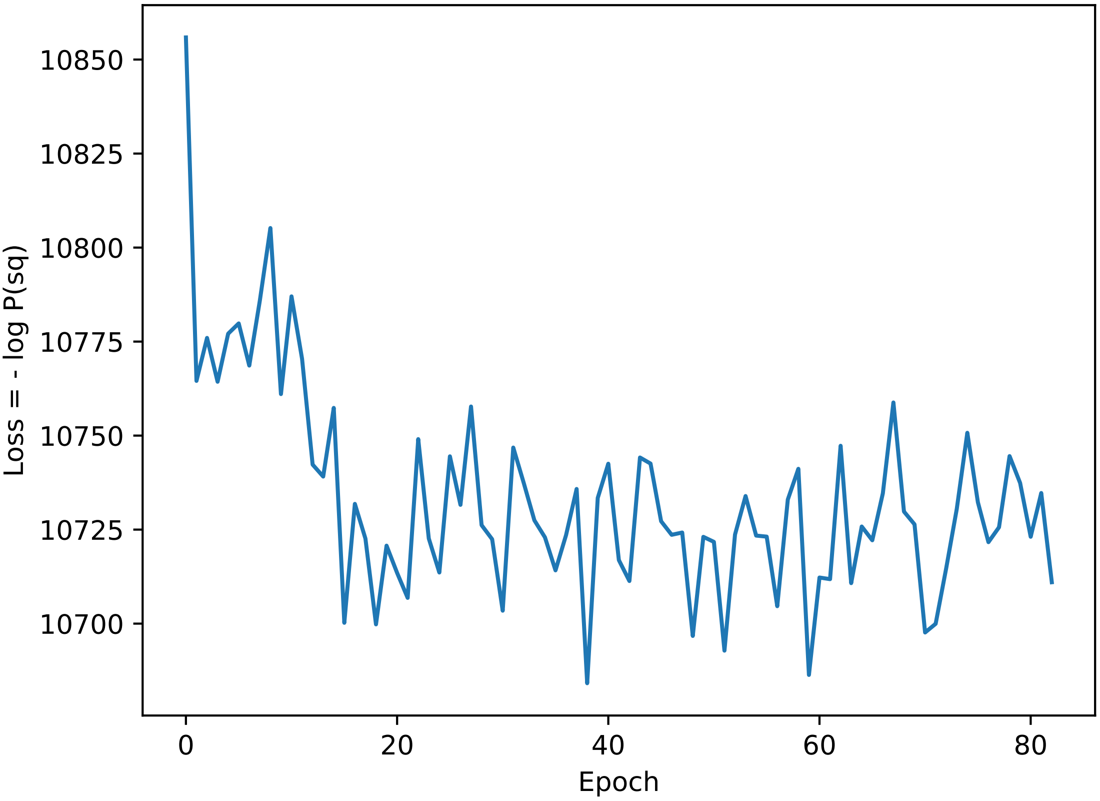

### losses_i94.pdf

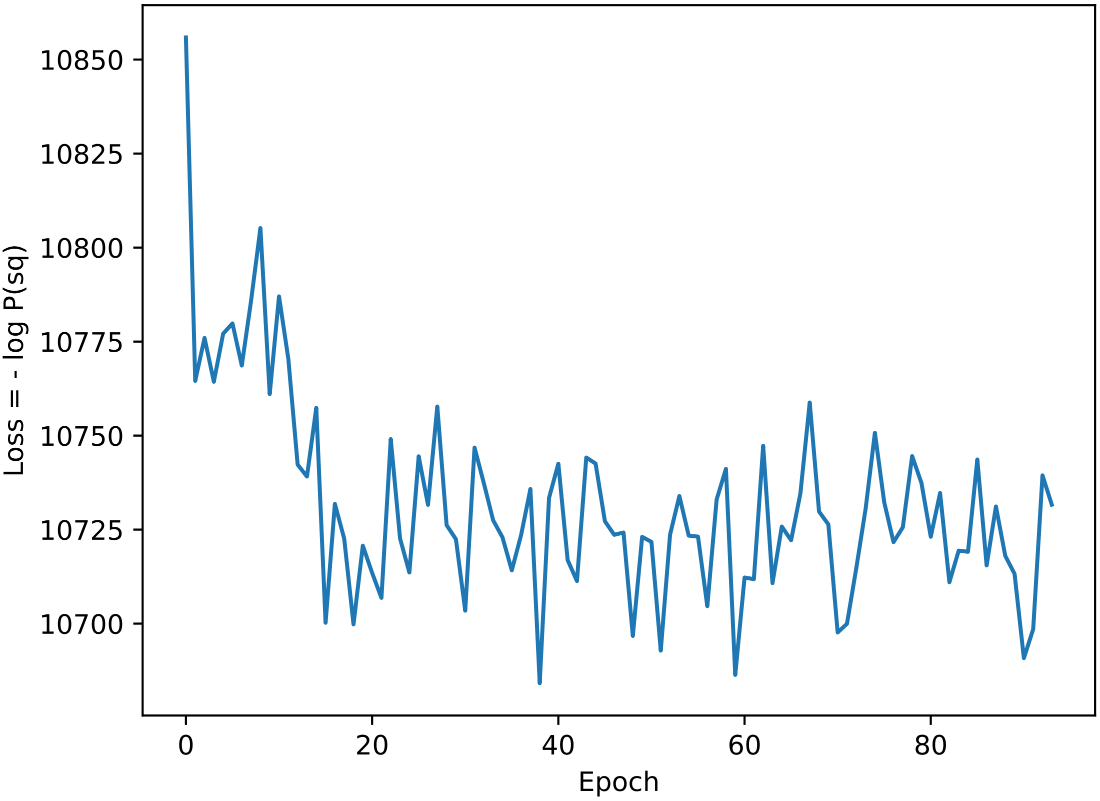

### losses_i95.pdf

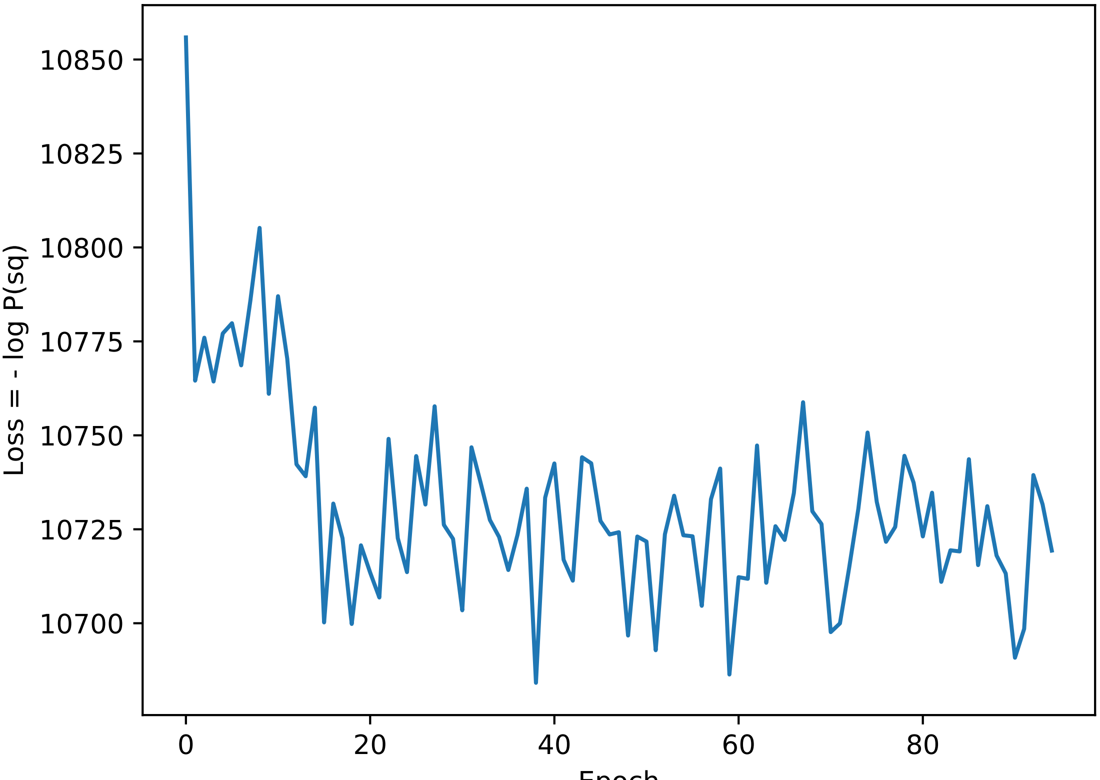

### losses_i96.pdf

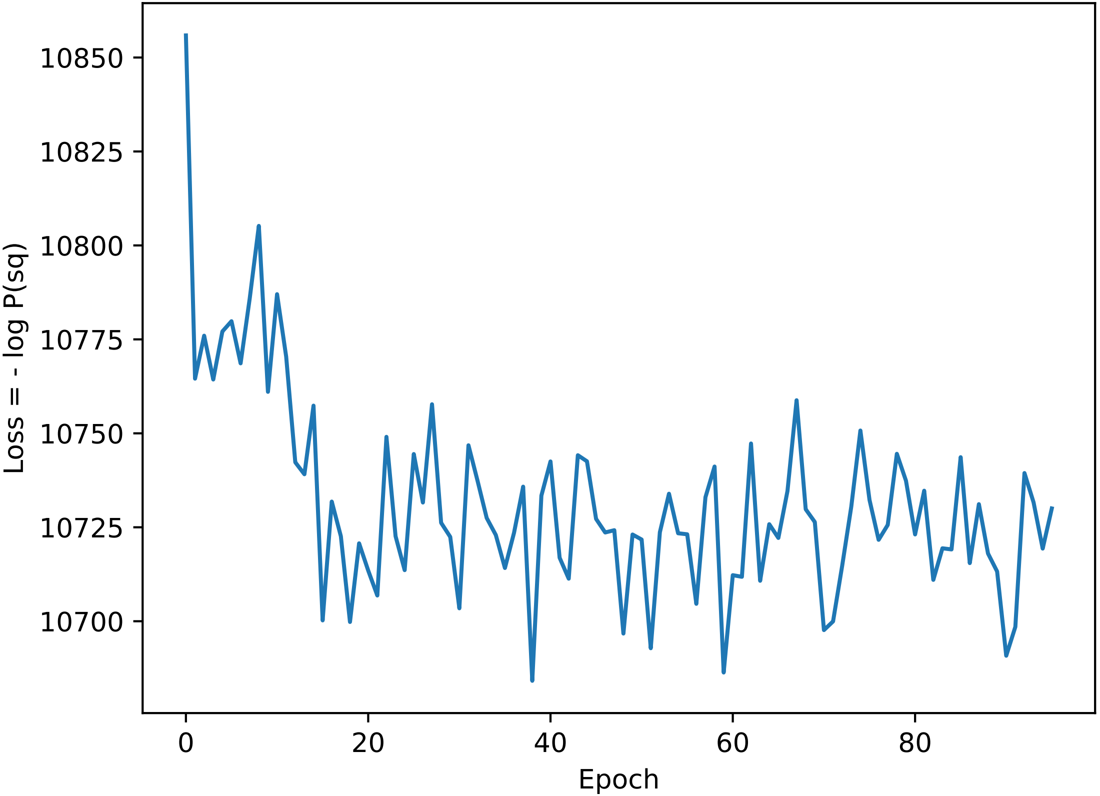

### losses_i97.pdf

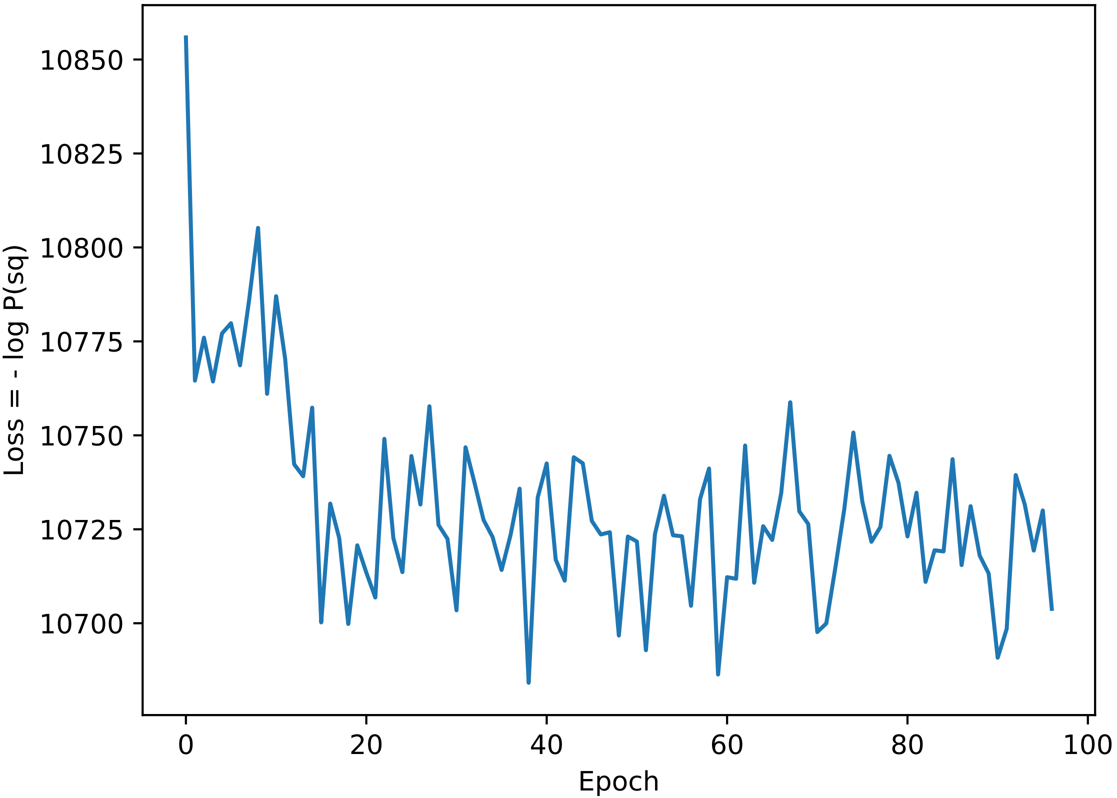
